# Intrinsic features of the RNase E membrane targeting sequence specify RNA degradosome organisation and activity

**DOI:** 10.64898/2026.03.25.714103

**Authors:** Sandra Amandine Marie Geslain, George Edward Allen, Johan Geiser, Peter Redder, Martina Valentini

## Abstract

In bacteria, transcription and RNA degradation are physically separated via segregation of the main ribonucleolytic machinery - the RNA degradosome - into phase-separated or membrane-anchored molecular assemblies driven by RNase E. Despite the widespread conservation of an amphipathic membrane anchor (MTS) in RNase E, the regulatory information embedded within this sequence and its biological importance remain poorly understood. Here, we have studied the importance of the *Pseudomonas aeruginosa* RNase E MTS for bacterial fitness or virulence and assessed its interchangeability. We show that amphipathicity is dispensable for foci scaffolding but necessary for proper foci morphology, dynamics, and localisation, although sequence modulates foci behaviour. Loss of the MTS additionally causes a drastic sensitivity to high salinity and a consistent virulence defect in *Galleria mellonella* larvae. Moreover, transcriptomics and analysis of mRNA spatial organisation reveal that the MTS mutant has specific stabilisation of localised membrane protein-encoding transcripts, together with abnormal operon processing. Altogether, our study highlights the elegant MTS-mediated control of spatial organisation and target selection, shaping the transcriptome and bacterial stress response.

## Introduction

Cells are uniquely organised and layered, a core property of life. Initially discovered in eukaryotic cells, intricate cytosolic organisation has also been observed in bacterial cells (Irastortza-Olaziregi & Amster-Choder, 2021). Notably, despite lacking membrane-bound organelles, bacteria are able to separate core cellular processes such as transcription, translation, or degradation by excluding or localising specific machineries to the cell pole, the midcell region, or the membrane (Irastortza-Olaziregi & Amster-Choder, 2021). Such organisation is largely driven by the nucleocytoplasmic ratio (NC ratio), a feature that shapes the biophysical properties of the cytoplasm (Gray *et al*, 2019). Bacteria with a high NC ratio, such as *Caulobacter crescentus*, exhibit a nucleoid that occupies most of the cytosol: ribosomes are mixed with the DNA mesh and diffusion rates are greatly decreased, causing most mRNAs to remain near their transcription site throughout most of their lifetime. On the other hand, in bacteria with a low NC ratio like *Escherichia coli*, the nucleoid is restricted to the middle of the cell and it excludes ribosomes, which are thus found enriched near the cell poles or the membrane (Gray *et al*., 2019; Irastortza-Olaziregi & Amster-Choder, 2021). In these bacteria, mRNAs are not enriched near their genomic loci and are instead found in various cellular locations throughout the entire cell, with most mRNAs coding for inner membrane proteins localising near the membrane (Kannaiah *et al*, 2019; Moffitt *et al*, 2016). The RNA degradosome, a key multiprotein complex involved in the processing of mRNAs, maturation of rRNA or tRNA, and in RNA decay, also exhibits a distinct subcellular localisation (Bandyra *et al*, 2013; Mackie, 2013; Tejada-Arranz *et al*, 2020). In *C. crescentus*, and in other species with a high NC ratio, the RNA degradosome assembles into large cytoplasmic foci termed bacterial ribonucleoprotein bodies (BR-bodies), organised by its core component RNase E (Al-Husini *et al*, 2018; Al-Husini *et al*, 2020; Gray *et al*., 2019). BR-bodies assembly is RNA-dependent and occurs through RNase E-mediated liquid-liquid phase separation (Al-Husini *et al*., 2018; Gray *et al*., 2019). In most Gammaproteobacteria, the RNA degradosome also localises as foci, but RNase E additionally harbours a conserved amphipathic helix of approximately 15 residues, called the membrane targeting sequence (MTS). In *E. coli*, it has been shown that the MTS positions the foci near the membrane (Khemici *et al*, 2008; Strahl *et al*, 2015) and an MTS-deleted *E. coli* RNase E harbours a diffuse subcellular localisation, indicating that the MTS is required for both membrane binding and foci assembly (Strahl *et al*., 2015; Thappeta *et al*, 2025). In addition, a recent study has shown that an *E. coli* RNase E mutant lacking the MTS exhibits a slower growth rate and accumulates high levels of 20S and 40S aberrant ribosomal particles, which the authors attribute to a premature cleavage of rRNA precursors resulting in accumulation of abnormally folded rRNA particles (Hadjeras *et al*, 2023). These results are consistent with a model in which separation of the RNA degradosome from the nucleoid avoids futile degradation of ribosome assembly intermediates, a reasoning that likely also applies to nascent mRNAs (Bandyra *et al*., 2013; Hadjeras *et al*., 2023; Mackie, 2013).

Several studies have shown that the interplay between mRNA localisation and RNA degradosome spatial organization influences mRNA half-life. Moffitt et al. (2016) demonstrated that, in *E. coli*, mRNAs encoding inner membrane proteins exhibit shorter half-lives than other transcripts, a difference which is abolished in a strain expressing an RNase E variant lacking the MTS (Moffitt *et al*., 2016). Notably, the MTS-deleted strain also displays a global increase in mRNA half-life, a phenotype independently reported by Hadjeras et al. (2019), consistent with earlier observations that delocalisation of RNase E from the membrane can impair its catalytic activity (Hadjeras *et al*, 2019; Murashko *et al*, 2012). Further supporting the importance of proper RNA degradosome localisation, a recent study showed that loss of MinD, which transiently interacts with RNase E via the MTS, leads to RNase E accumulation at the cell poles, resulting in disrupted polar mRNA localisation and reduced half-lives of polar transcripts (Kannaiah *et al*., 2024).

Despite the growing evidence about the need to compartmentalise the RNA degradosome, how this separation contributes to bacteria fitness and stress adaptation remains largely unknown. To our knowledge, the phenotypic effect of RNA degradosome membrane delocalisation reported to date is a mild growth defect in rich medium (Hadjeras *et al*., 2023). In addition, the MTS sequence is known to be rather conserved in all Gammaproteobacteria, but its recently described regulatory interaction with MinD in *E. coli* raises an interesting question: is the MTS involved in additional roles beyond membrane anchoring, such as fine control of RNA degradosome clustering and activity?

To bridge this gap, we studied membrane anchoring of the RNA degradosome in the WHO high-priority pathogen *Pseudomonas aeruginosa*, a bacterium especially suited to RNA metabolism studies due to its adaptability and intricate regulatory networks (Galán-Vásquez *et al*, 2011). By comparing a mutant lacking the RNase E MTS with a chimeric RNase E mutant bearing a heterologous amphipathic helix, we have assessed the interchangeability and in vivo importance of the RNase E MTS. Overall, our results show that MTS-mediated anchoring is dispensable for assembly of RNA degradosome foci but is required for proper foci dynamics and subcellular localisation. Importantly, deletion of the MTS impairs adaptation to specific stresses and virulence in a *G. mellonella* model. Furthermore, we uncover a previously unrecognized role for the MTS in RNase E operon-specific mRNA processing, affecting a subset of transcripts encoding inner membrane proteins. Overall, our work establishes a novel role for the RNase E MTS in mRNA target selection and in normal adaptative stress response, likely sustained by the elegant process of MTS-mediated RNA degradosome clustering and dissolution.

## Results

### RNase E MTS deletion redistributes RNA degradosome foci in *P. aeruginosa* and is complemented by MinDα

In Beta and Gammaproteobacteria, most RNase E homologs harbour a membrane-targeting sequence (MTS) defined by a conserved amphipathicity (Aït-Bara *et al*, 2015; Bandyra *et al*., 2013; Khemici *et al*., 2008). We previously confirmed the presence of an MTS within *P. aeruginosa* RNase E (PaRNase E) and showed that the protein forms discrete foci at the membrane (Geslain *et al*, 2025). To investigate the subcellular localisation of PaRNase E upon loss of the MTS, we constructed a strain expressing PaRNase E^ΔMTS^-msfGFP from the native chromosomal locus. Visualisation of the signal revealed that the PaRNase E^ΔMTS^-msfGFP is still capable of assembling foci (Fig. 1A); however, these foci have statistically significant higher fluorescence intensity values and rounder outlines than the ones formed by the full-length PaRNase E (Fig. 1B). Additionally, the average subcellular localisation of PaRNase E^ΔMTS^-msfGFP is shifted compared to the native protein: although foci still tend to be enriched near the cell periphery, we observe a striking increase in foci density near the core region of the cells and a significantly higher frequency of foci located at the cell poles (12,4 % vs 5,7 %) and midcell (10% vs 7,3%) compared to foci formed by the full-length PaRNase E (Fig. 1C). To verify that the foci observed in the PaRNase E^ΔMTS^-msfGFP were RNA-dependent assemblies and not caused by protein aggregation or misfolding, we treated the cells with rifampicin to deplete cellular RNA due to transcription inhibition. Visualisation of the cells after 30 minutes showed complete dissolution of PaRNase E^ΔMTS^-msfGFP foci as observed with the full-length PaRNase E (Fig. 1D). Additionally, we observed a completely homogeneous subcellular localisation of rifampicin-treated PaRNase E^ΔMTS^-msfGFP, in contrast to the membrane-enriched peripheral localization of PaRNase E full-length (Fig. 1D), which further confirms that deletion of the MTS delocalises PaRNase E, and thereby the RNA degradosome, from the membrane.

**Figure 1:**
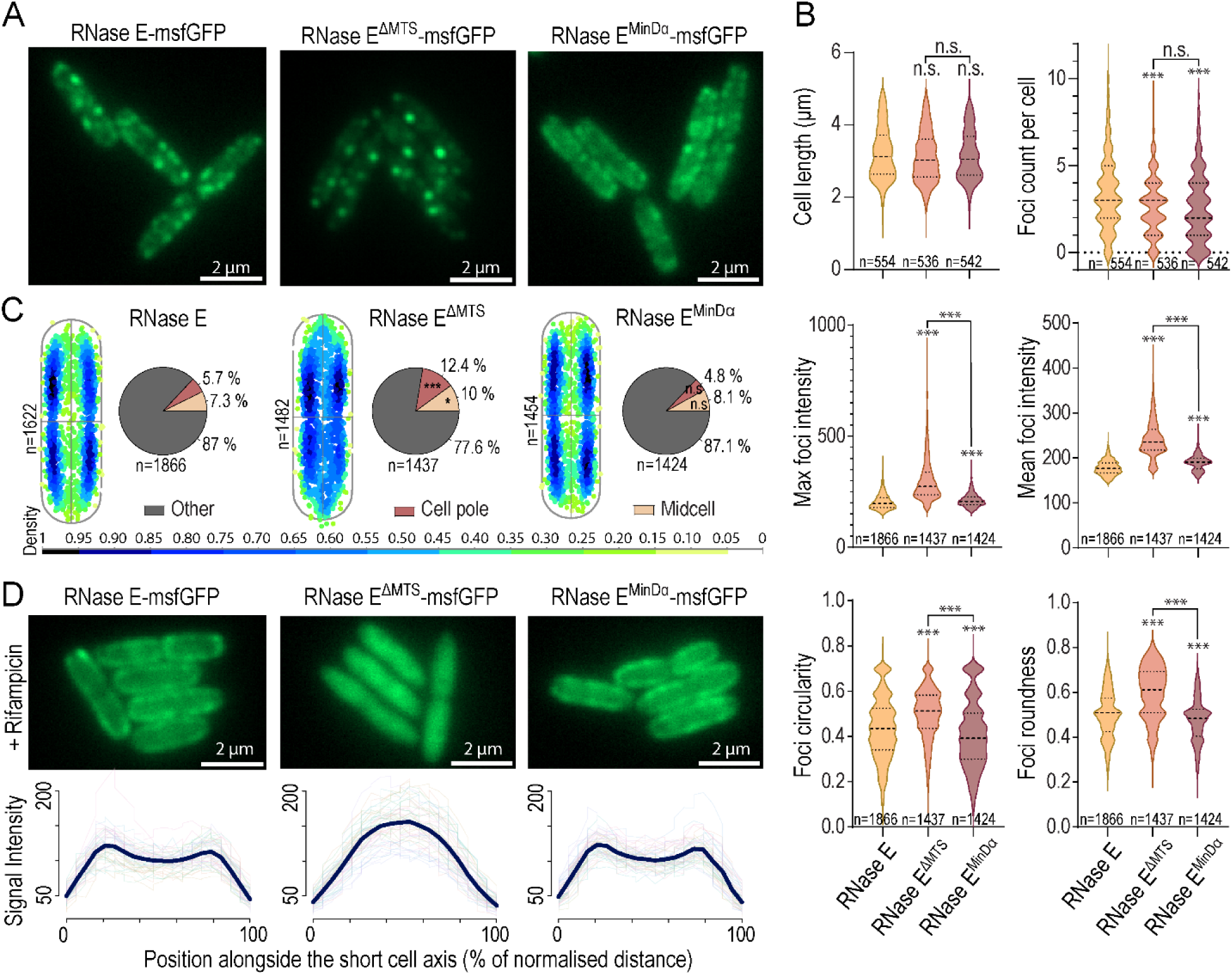
Deletion of the RNase E MTS causes foci morphology and localisation defects which can be complemented by reinsertion of the unrelated MinDα. **(A)** Representative images of RNase E, RNase E^ΔMTS^, or RNase E^MinDα^ C-terminally tagged with msfGFP (chromosomal expression from the native locus). **(B)** Violin plots (with median and quartiles) showing the quantification of cell length, RNase E foci count per cell, foci mean or max intensity and foci circularity or roundness from images of msfGFP-tagged RNase E variants (as described) acquired in three independent biological replicates. The analyses were performed using the MicrobeJ plugin of Fiji as described in Materials and Methods. An unpaired t-test with Welch’s correction was done using GraphPad to compare mutant RNase E variants to the native RNase E (default comparison) or to each other (highlighted by brackets). n.s.: not significant (99% confidence interval); ***: p-val < 0.000001. **(C)** XY cell density graph of the average distribution of RNase E foci in a bacterial cell (left) and pie charts (right) highlighting the proportions of RNase E foci found at the midcell, the cell poles, or elsewhere (“Other”). Localisation of the foci was automatically detected using the MicrobeJ plugin. The XY cell density plots were generated using MicrobeJ whereby each detected foci, represented by a dot, is coloured based on its vicinity with neighbouring foci. For details about the settings used, see Materials and Methods. Two-sample χ² tests for equality of proportions with continuity correction were carried out. *: p-val < 0.01; ***: p-val < 0.000001. **(D)** Representative images (top panels) of RNase E, RNase E^ΔMTS^, or RNase E^MinDα^ C-terminally tagged with msfGFP (chromosomal expression from the native locus) after 30 minutes exposure to Rifampicin (100 μg/mL) and average fluorescence intensity profiles (bottom panels) from the short cell axis of the Rifampicin-exposed strains.

Although the primary sequence of the MTS varies across species, conserved hydrophobic or polar residues at defined positions maintain its amphipathic nature (Aït-Bara *et al*., 2015; Appendix Fig. S1). To test whether this physicochemical property is sufficient for function, we constructed a chimera in which the amphipathic helix of the unrelated *E. coli* MinD protein (MinDα, Appendix Fig. S1), was chromosomally inserted back at the native position in our strain expressing PaRNase E^ΔMTS^-msfGFP (Szeto *et al*, 2003; Zhou & Lutkenhaus, 2003). Visualisation of this strain expressing PaRNase E^MinDα^-msfGFP and automated quantification of foci morphology and localisation revealed a nearly complete complementation to a WT-like appearance (Fig. 1A-C). Indeed, the average foci subcellular localisation profile and counts at midcell or cell poles are virtually identical to those of full-length PaRNase E-msfGFP, while the foci are significantly less bright and less round than those of PaRNase E^ΔMTS^-msfGFP. Exposure of the strain expressing PaRNase E^MinDα^ -msfGFP to rifampicin also results in a peripheral localization, with signal intensity profiles virtually identical to those of the full-length RNase E (Fig. 1D). Interestingly, we observe that the PaRNase E^MinDα^ foci are visually less well-defined, or more diffused, than those of the full-length RNase E (Fig 1A). Quantification of foci morphology confirms these observations and reveals significant differences in foci intensity and circularity/roundness compared to the full-length RNase E (Fig. 1B), suggesting that intrinsic features of the amphipathic helix subtly regulate RNA degradosome clustering.

### PaRNase E MTS deletion causes stalling of the RNA degradosome foci and is partially rescued by MinDα

In *E. coli* and *C. crescentus*, RNase E foci are known to dynamically form and dissolve within the range of seconds (Al-Husini *et al*., 2018; Strahl *et al*., 2015). The altered morphology and increased intensity of PaRNase E^ΔMTS^ foci prompted us to investigate whether their assembly or dynamics were affected.

To rule out an impaired recruitment of the RNA degradosome components PNPase and RhlB, we constructed a *P. aeruginosa* strain carrying both RhlB-msfGFP and PNPase-mTagBFP2 fusions at their native gene loci, together with a mCherry-tagged variant of PaRNase E (full-length, RNase E^ΔMTS^, or RNase E^MinDα^). Using structured illumination microscopy (SIM), we observed a strong colocalisation of the three protein partners irrespective of the co-expressed PaRNase E variant (Fig. 2), confirming that deletion of the MTS or its replacement with the MinDα do not prevent recruitment of PNPase and RhlB into the RNase E-driven foci.

**Figure 2:**
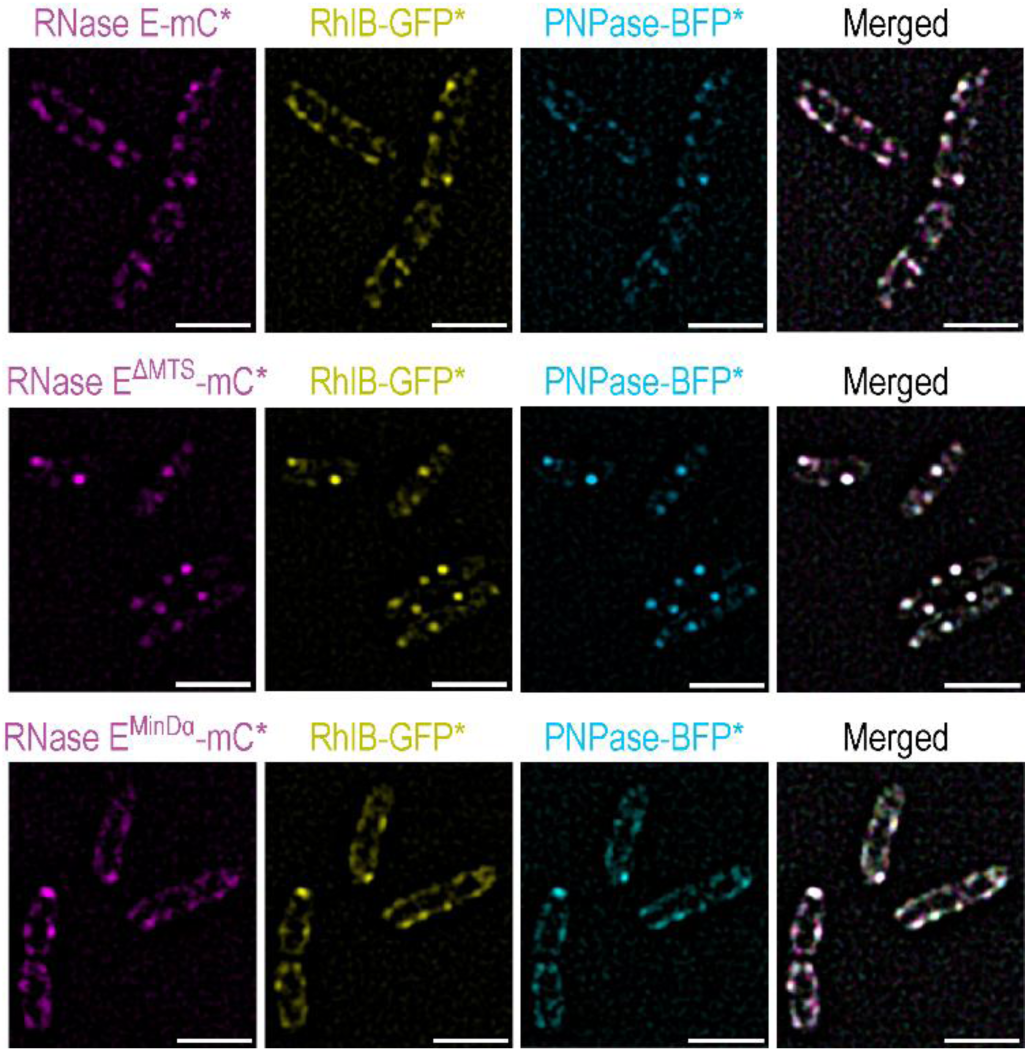
Deletion of the MTS or its replacement with MinDα do not impact recruitment of RhlB and PNPase into the RNase E-driven foci Visualisation with structured illumination microscopy (Zeiss ELYRA7) of native or mutated RNase E-mCherry fusions as described (left column), RhlB-msfGFP fusion (middle column), or PNPase-mTagBFP2 fusion (second to last column) co-expressed in the same strain. mC*: mCherry; *GFP: msfGFP; BFP*: mTagBFP2. The three channels were fused using Fiji to highlight signal overlap (Merged images).

We next used time-lapse SIM to investigate whether foci dynamics were altered in the PaRNase E mutants. Strikingly, foci formed by PaRNase E^ΔMTS^ were severely stalled, with most remaining assembled and stationary throughout the 120-second acquisition period (Fig. 3B and Supp. Movie 2). This is in sharp contrast with foci assembled by PaRNase E full-length, which seem to rapidly dissolve and reassemble within the 5-seconds imaging frames, giving the impression that PaRNase E is circling through the bacterial membrane (Fig. 3A and Supp. Movie 1). Interestingly, we observed that the PaRNase E^MinDα^ foci seem to have an intermediate behaviour: while most foci are dynamic and rapidly disappear and reform within 5 seconds, as observed with the full-length PaRNase E, some cells harbour several larger foci that remain stalled throughout the course of the time-lapse acquisition period (Fig. 3C and Supp. Movie 3).

**Figure 3:**
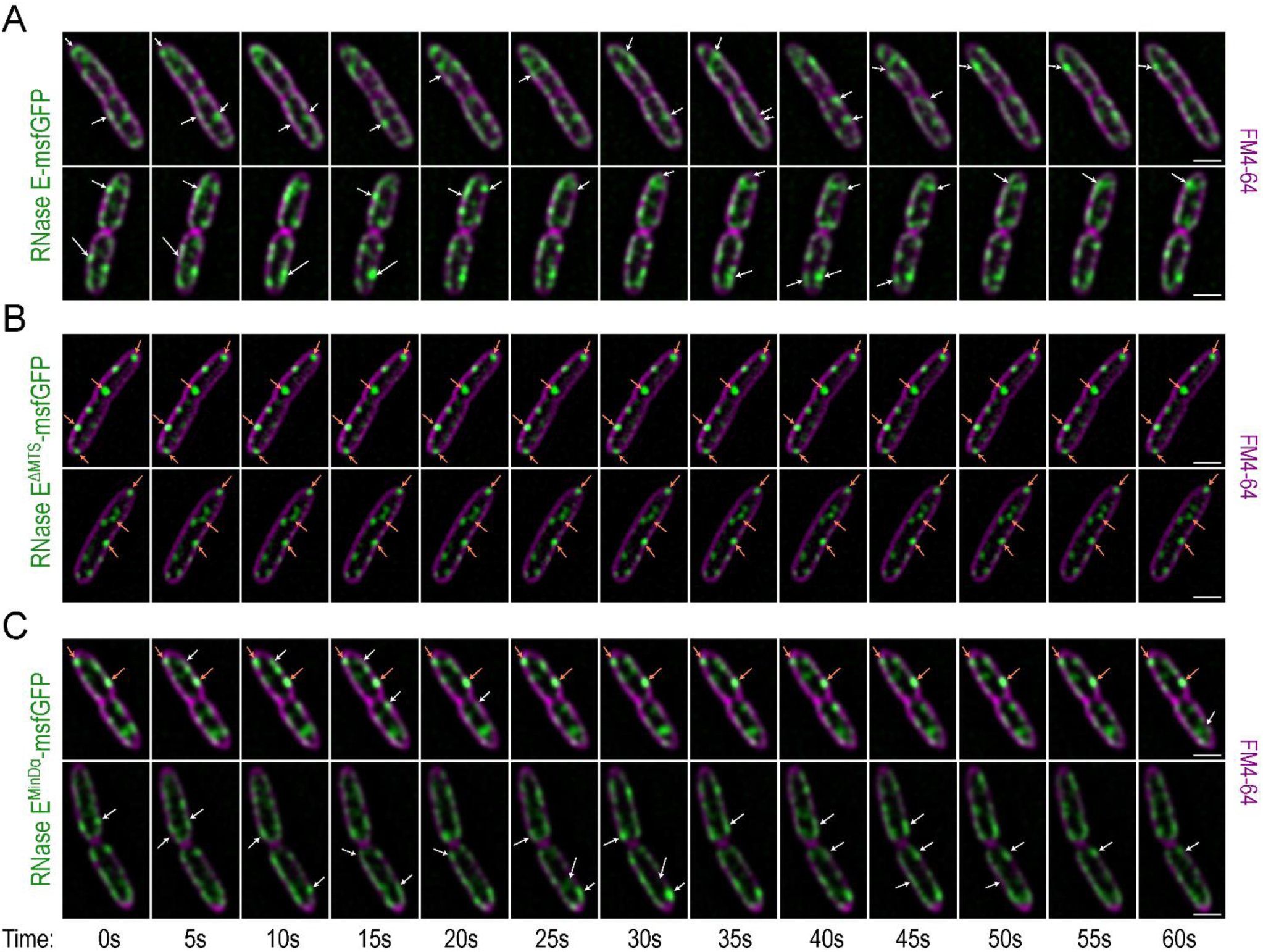
Loss of the MTS results in an immobilisation of the RNase E foci which can be rescued by reinsertion of MinDα. SIM Time-lapse visualisation of live bacterial cells chromosomally expressing **(A)** RNase E-msfGFP, **(B)** RNase E^ΔMTS^-msfGFP, or **(C)** RNase E^MinDα^-msfGFP. Membranes were stained using the FM4-64 dye. Foci that visually form, dissolve, or diffuse laterally are highlighted by white arrows, whereas foci that remain static throughout the acquisition are indicated by orange arrows. Scale is 1 μm. Channels showing the msfGFP and FM4-64 signal were merged for display using Fiji.

Overall, these results suggest that the amphipathic nature of the MTS drives RNA degradosome localization and dynamics, whereas its specific sequence modulates foci behaviour.

### The PaRNase E MTS is required for specific stress adaptation and can be substituted by MinDα

Despite the widespread conservation of the MTS in RNase E homologs, its functional importance for bacterial physiology and adaptive capacity remains unclear (Hadjeras *et al*., 2023; Hadjeras *et al*., 2019). To address this, we performed an extensive phenotypic characterization of the *rne*ΔMTS mutant, testing for growth on minimal medium or under acidic, basic, or ethanol stress conditions, as well as growth under exposure to detergents, salts, high temperatures, or iron chelating agents (Fig. 4 and Fig. EV1). While the mutant grew nearly identically to the WT strain at 37°C, with only minor growth defects observed at 16°C, it displayed a pronounced and specific growth defect in rich medium supplemented with 0.75 M NaCl or KCl, or 0.5 M (NH_4_)_2_SO_4_ (Fig. 4A and Fig. EV1). These defects were not attributable to osmotic stress alone: the *rne*ΔMTS mutant grew comparably to WT in medium supplemented with 0.75 M sucrose (Fig. 4A). Together, these results indicate that deletion of the MTS confers a specific sensitivity to elevated ionic strength rather than to increased osmolarity per se. To assess virulence, we performed a killing assay using *G. mellonella* larvae, an infection model highly susceptible to *P. aeruginosa* infections (Beeton *et al*, 2015). Although near-complete mortality was observed by 24 h post-infection (n = 77 larvae), we detected a statistically significant delay of one to two hours in killing time, indicating that the *rne*ΔMTS mutant is partially compromised in its ability to establish or sustain infection (Fig. 4B). Of note, total levels of full-length RNase E and RNase E^ΔMTS^ were similar in exponentially growing cultures, ruling out altered protein abundance as the cause of the observed phenotypes (Appendix Fig. S2).

**Figure 4:**
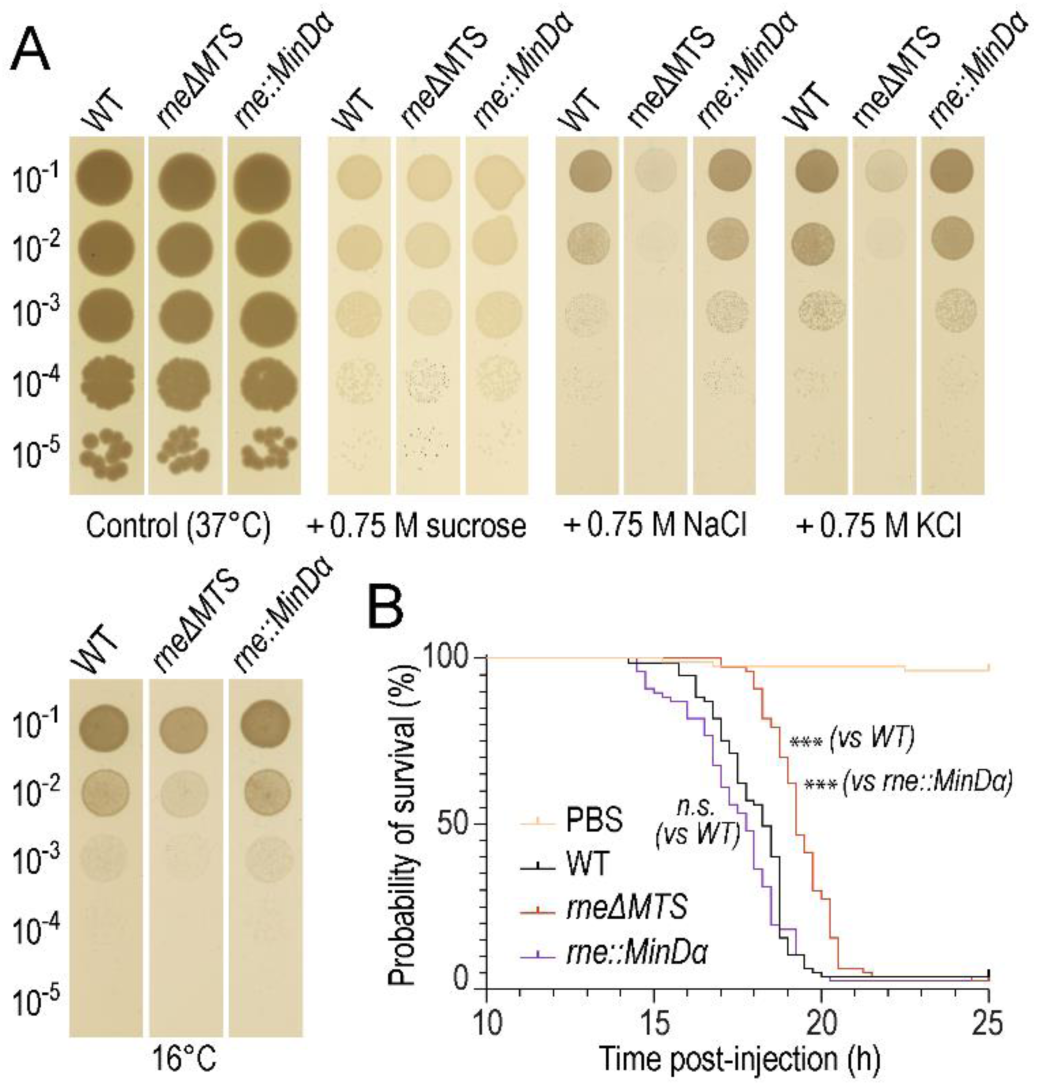
Reinsertion of MinDα can fully complement the phenotypic defects of the *rne*ΔMTS strain in high salinity conditions or during infection of *Galleria mellonella*. **(A)** Growth of PAO1 WT, *rne*ΔMTS, or *rne*::MinDα on solid medium at 37°C under control conditions or with addition of 0.75M NaCl, KCl or sucrose to the growth medium. Growth at 16°C was also tested (lower panel). Three independent biological replicates were done; a representative plate is shown. **(B)** Killing curves of *G. mellonella* larvae after injection with 10 μL of PAO1 WT, *rne*ΔMTS, or *rne*::MinDα dilutions. The control group was injected with 10 μL of sterile PBS instead. Death was monitored over 14-25 hours post-injection. Biological triplicates were done using 20-30 larvae per replicate (total of 77-80 larvae per strain tested). Statistical significance was assessed by performing a Kaplan-Meier survival analysis in GraphPad Prism using a log-rank (Mantel-Cox) test and comparing the mutant RNase E variants to the native RNase E or to each other as described. ***: p-value <0.000001; n.s.: not significant (99% confidence interval).

To assess whether the observed defects of the *rne*ΔMTS strain could be rescued by the *E. coli* MinDα, we reinserted MinDα back into the *rne*ΔMTS strain (at the native chromosomal position). Strikingly, the *rne*::MinDα strain can grow under stress or infect *G. mellonella* larvae virtually identically to the WT strain and is thus a full complementation of the *rne*ΔMTS defects (Fig. 4A-B). These results highlight the importance of the amphipathic nature of the helix for proper RNase E and RNA degradosome function under challenging conditions such as high salinity or host colonization.

### Deletion of PaRNase E MTS specifically stabilises a subset of transcripts encoding inner membrane proteins and impacts differential processing of some polycistronic transcripts

We previously performed transcriptomic analyses of two RNase E mutants lacking the C-terminal scaffolding domain, one mutant additionally lacking the MTS and the other retaining it, and found that despite a comparable number of misregulated transcripts (400-450 transcripts in total), the affected gene sets showed little overlap (Geslain *et al*., 2025). This divergence prompted us to analyse the transcriptome of the *rne*ΔMTS mutant to clarify the specific contribution of the MTS to gene regulation. Triplicate total RNA samples were first analysed by capillary electrophoresis, which revealed no difference in size profiles compared to WT, contrasting with previous reports in *E. coli* (Appendix Fig. S3) (Hadjeras *et al*., 2023). Differential expression analysis identified 93 significantly misregulated genes only (Fig. 5A), indicating a limited transcriptional impact compared to the broader changes previously observed in the two aforementioned RNase E mutants. Gene Ontology (GO) enrichment analysis of the differentially regulated transcripts revealed overrepresentation of genes encoding transmembrane transporters as well as proteins involved in secondary metabolite biosynthesis or protein folding and proteolysis (Fig. 5B). A previous study reported preferential stabilization of transcripts encoding inner membrane proteins in the *E. coli rne*ΔMTS strain (Moffitt *et al*., 2016). We therefore examined the predicted subcellular localisation of the proteins encoded by the misregulated transcripts using published annotations of secretion and transmembrane signals in *P. aeruginosa* (Lewenza *et al*, 2005). Among the 54 upregulated transcripts, genes encoding proteins containing a predicted transmembrane helix were significantly enriched (n=23), whereas transcripts lacking predicted localisation signals were depleted (n=16), both relative to genome-wide proportions and to the set of downregulated transcripts (Fig. 5C). These results confirm that deletion of the MTS preferentially affects a subset of transcripts encoding inner membrane proteins instead of having a pleiotropic effect on the transcriptome. To investigate whether RNase E cleavage sites were affected, we have optimised and performed a variant of 5’ monophosphorylated RNA sequencing which we named GC-EMOTE (Redder, 2018). Scatter plot comparisons of unique 5’ ends sequenced in the *rne*ΔMTS or WT reveal a similar global profile, although a subset of 5’ ends appear to be significantly depleted or enriched in the *rne*ΔMTS strain (Fig. 5D). To assess whether the sequenced 5’ ends were likely to have been generated by RNase E, we separately analysed 5’ ends with dinucleotide sequences either most or least compatible with known RNase E consensus sequence. Our analyses reveal that the profile of the most RNase E-like 5’ ends has a broader distribution compared to the least RNase E-like 5’ ends, consistent with the idea that RNase E cleavage site selection is partially affected in the *rne*ΔMTS mutant (Fig. 5D). To view how differential processing activity may contribute to the observed gene misregulation, we have analysed the RNA-seq and GC-EMOTE coverage of operons affected in our transcriptome analysis. Strikingly, genes found within the same polycistronic transcripts can have steady-state levels that are differentially altered in the *rne*ΔMTS mutant. For example, although arcD or atpI are respectively downregulated or upregulated in the *rne*ΔMTS mutant, most other genes in these operons follow the opposite pattern. These alterations correlate with a GC-EMOTE peak profile showing different peak depth between the two strains, suggesting that differential RNase E-mediated cleavage site selection accounts for the changes in polycistronic mRNA levels observed in the *rne*ΔMTS.

**Figure 5:**
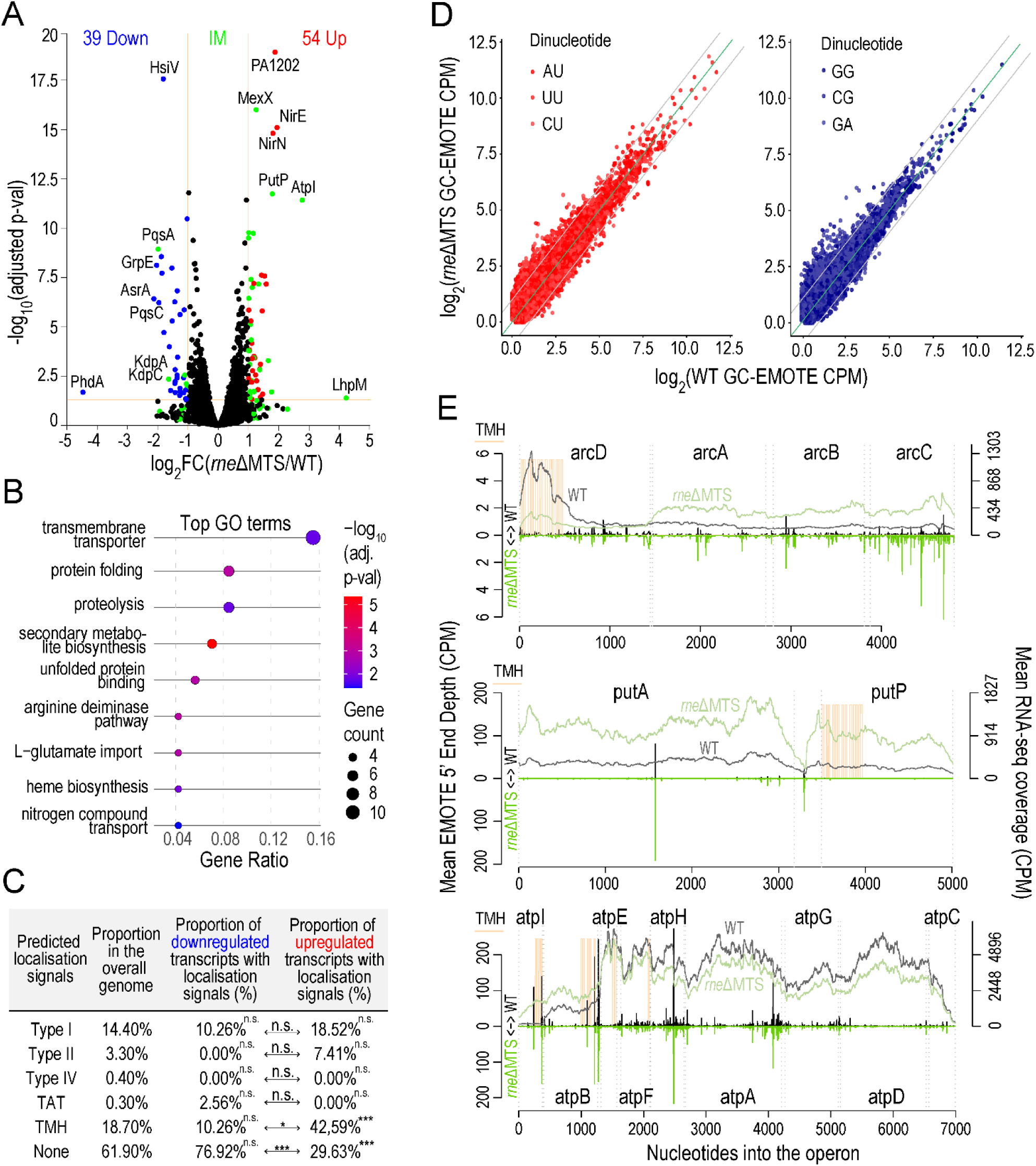
RNA-seq and GC-EMOTE analyses highlight the importance of the MTS for expression levels and processing of a subset of operons and transcripts encoding inner membrane proteins **(A)** Volcano plots highlighting genes upregulated (red) or downregulated (blue) in the *rne*ΔMTS mutant compared to the WT strain. Genes coding for inner membrane (IM) proteins are coloured in green. Annotations of inner membrane proteins were retrieved from the Pseudomonas.com database. **(B)** Gene Ontology (GO) analysis of the RNA-seq dataset highlighting overrepresented gene functions misregulated in the *rne*ΔMTS mutant. **(C)** Table highlighting the proportion of genes containing various localisation signals in the sets of downregulated or upregulated transcripts compared to the overall genome. A two-sample χ² test for equality of proportions with continuity correction was used to compare the proportions from each set of transcripts to the overall genome proportions (default comparison, indicated next to each value) or to each other (indicated by arrows). n.s.: not significant; *: p-val < 0.01; ***: p-val < 0.0001. **(D)** Scatter plots of GC-EMOTE unique monophosphorylated 5’ ends in the WT strain (x axis, counts per million) or *rne*ΔMTS strain (y axis, counts per million) bearing either the most RNase E-compatible AU, UU, or CU dinucleotides (red graph, left) or the most RNase E-incompatible GG, CG, or GA dinucleotides (blue graph, right) at the cleavage site position. Dinucleotide compatibility with RNase E cleavage was assessed by generating a score based on the RNase E consensus logo plot sequence published in (Chao *et al*, 2017). Nucleotide weights (2^bits) for positions one and two were calculated from their graph of information content (bits) and multiplied to give the dinucleotide score. **(E)** GC-EMOTE peaks and RNA-seq coverage of three operons containing at least one transcript significantly misregulated in the *rne*ΔMTS transcriptomics analysis (panel A). The positions of transmembrane helices in the encoded protein are highlighted in light orange. RNA-seq coverage shows differential transcript stabilisation within the arc and the atp operons in the *rne*ΔMTS, correlating with a different overall peak profile in the GC-EMOTE dataset.

### Investigation of mRNA subcellular localisation by smFISH reveals a specific membrane localisation of a model TMH-encoding transcript stabilised in the *rne*ΔMTS mutant

Because *E. coli* monocistronic or polycistronic mRNAs that encode for at least one inner membrane proteins are known to themselves locate near the inner membrane (Kannaiah *et al*., 2019; Moffitt *et al*., 2016; Nevo-Dinur *et al*, 2011), we hypothesized that most of the TMH-encoding transcripts which are upregulated in the *rne*ΔMTS genetic background become shielded from the RNA degradosome due to their segregation at the inner membrane. Choosing three well-expressed transcripts showing different patterns in the RNA-seq, namely asrA (cytosolic protein, downregulated), pqsAB (inner membrane protein for PqsA, both downregulated) and putP (inner membrane protein, upregulated), we specifically investigated mRNA spatial subcellular localisation in *P. aeruginosa* using non-multiplexed single molecule RNA Fluorescence In Situ Hybridisation (smFISH).

Epifluorescence imaging of smFISH-treated bacterial samples revealed discrete fluorescent foci in both the WT and *rne*ΔMTS strains for all three genes tested (Fig. 6A and Fig. EV2). In smFISH, a single mRNA molecule typically appears as one bright, diffraction-limited focus due to signal amplification from the annealing of approximately 48 fluorescent probes per transcript (Skinner *et al*, 2013). To verify probe specificity, we generated deletion mutants for each of the three genes and confirmed that the signal observed in the WT and *rne*ΔMTS strains was specific to the target mRNAs (Fig. 6A and Fig. EV2). Automated quantification of the smFISH images showed that differences in foci number and intensity between WT and *rne*ΔMTS mirrored the RNA-seq results. Indeed, the upregulated putP transcript displayed increased foci counts and higher signal intensity in the *rne*ΔMTS strain, whereas the downregulated asrA and pqsAB transcripts exhibited reduced foci numbers and intensities compared to the WT (Fig. 6C–D). Although a few foci were automatically detected in the deletion mutants, their fluorescence intensity was minimal and consistent with background signal, and no foci could be visually observed.

**Figure 6:**
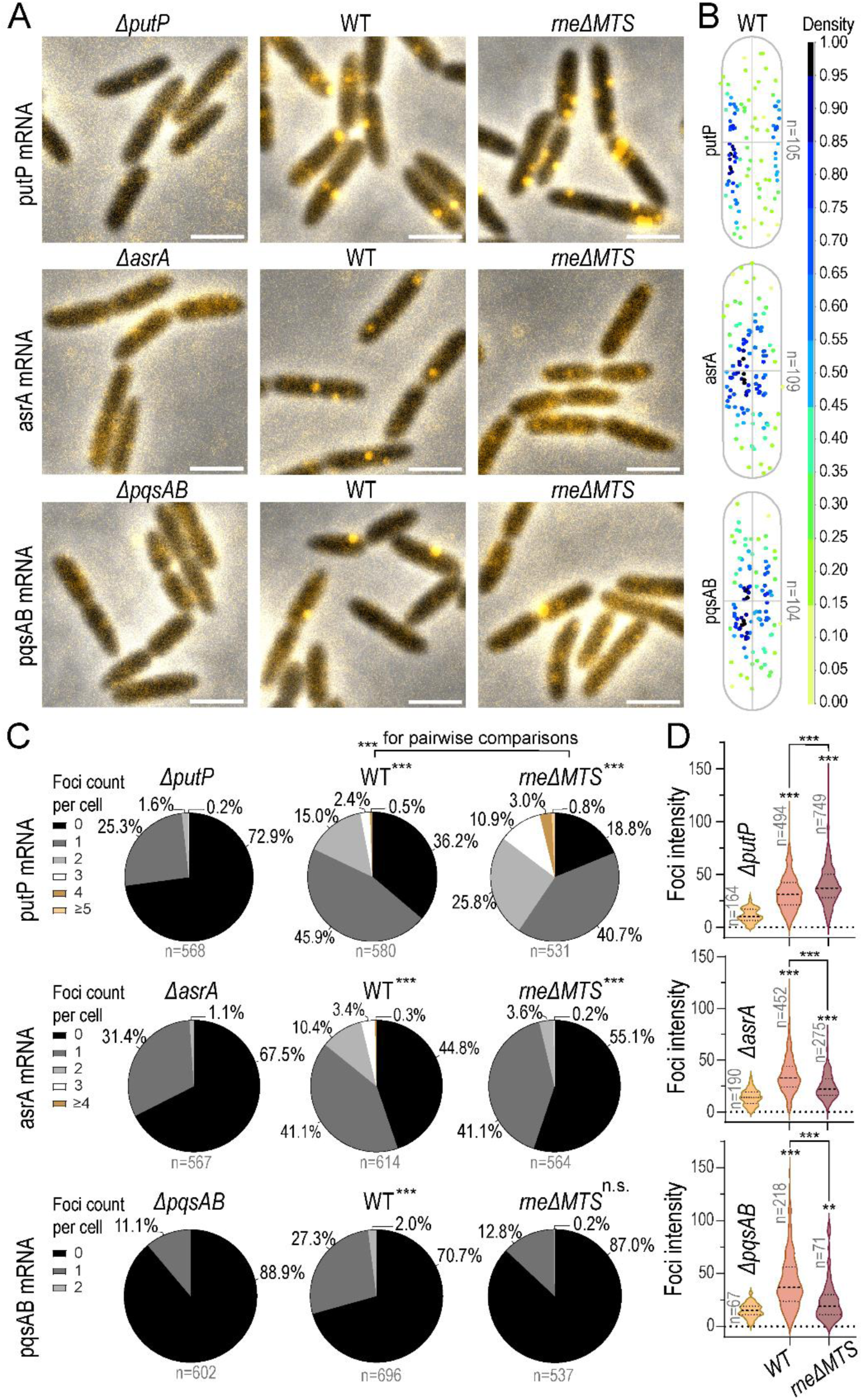
Fluorescence in situ hybridisation (FISH) reveals different overall localisation patterns of the putP, pqsAB, and asrA transcripts and confirms altered expression levels in the *rneΔMTS* strain. **(A-B)** Representative images (A) and XY cell density graph of the average distribution of FISH foci (B) for the putP (upper panels), asrA (middle panels) or pqsAB (lower panels) mRNAs. One foci theoretically corresponds to one mRNA molecule. Scale is 2 μm. The XY cell density graph of the average distribution of FISH foci was generated using MicrobeJ using three images for each transcript, each acquired in independent biological replicates. **(C-D)** Pie charts (C) and violin plots with median and quartiles (D) highlighting the proportions of cells having 0, 1, or more foci of putP (upper panels), asrA (middle panels) or pqsAB (lower panels) mRNA and the measured intensity of those foci, respectively. The foci detection and intensity measurement were automatically done with the MicrobeJ plugin in Fiji using images acquired in three independent biological replicates. One to three images were analysed for each biological replicate (approximately 125-250 cells per biological replicate). A two-sample χ² test for equality of proportions with continuity correction was used to compare the proportions of foci per cell (C) in the WT or *rne*ΔMTS strain relative to the negative control (default comparison) or relative to one another (highlighted by brackets). ***: p-value < 0.0001; n.s.: not significant. Unpaired t-tests with Welch correction were performed using GraphPad to compare the foci intensities (D) observed in the WT or *rne*ΔMTS strain relative to the negative control (default comparison) or relative to one another (highlighted by brackets). ***: p-value < 0.000001; **: p-value < 0.001.

We next investigated the subcellular distribution of putP, asrA, and pqsAB mRNA foci using stringent detection parameters (Fig. 6B). The distribution profiles reveal that putP mRNAs are preferentially enriched near the membrane, consistent with the localisation of the encoded inner membrane protein and with its stabilisation in the *rne*ΔMTS strain. Conversely, asrA mRNAs, which encode a cytosolic protein, are distributed throughout the cytoplasm but appear enriched towards the centre of the cell. Although individual pqsAB mRNA foci were observed near the membrane, the overall distribution profile suggests a ubiquitous localisation, with signal also concentrated near the centre of the cell.

Despite the inner membrane localisation of the PqsA protein, the observed pqsAB mRNA distribution indicates limited membrane enrichment, consistent with the downregulation of the pqsABCDE operon in the *rne*ΔMTS strain. Alternatively, processing of the pqsABCDE polycistronic mRNA may generate truncated transcripts of varying length showing distinct localization patterns, potentially accounting for the mixed distribution observed.

Overall, our smFISH images reveal distinct subcellular localization patterns for putP, asrA, and pqsAB mRNAs and provide independent support for their consistent differential regulation in the *rne*ΔMTS strain, highlighting an intricate connection between mRNA localisation and RNA degradosome segregation.

## Discussion

The interplay between RNA degradosome compartmentation and mRNA localisation has recently emerged as a core element influencing gene expression and mRNA stability (Irastortza-Olaziregi & Amster-Choder, 2021; Kannaiah *et al*, 2024; Moffitt *et al*., 2016). However, our understanding of the means through which bacteria spatially organise the RNA degradosome, as well as the biological consequences of this compartmentation, remains limited to a few case studies.

One key feature which determines the subcellular localisation of Gammaproteobacterial RNA degradosomes is the conserved RNase E membrane-targeting sequence (MTS). In the present study, we show that a *P. aeruginosa* RNase E mutant lacking the MTS is partially delocalised from the membrane. In the *rne*ΔMTS mutant, a broader RNase E subcellular distribution compared to the WT is observed, with a concomitant increase in the fraction of foci localising at the cell poles or at the midcell. Importantly, in the *rne*ΔMTS mutant, the RNA degradosome foci are still assembled but they have abnormal characteristics: foci are significantly larger, more rounded, and have an increased signal intensity compared to WT foci. Time-lapse SIM revealed that these foci are also severely stalled and cannot rapidly dissolve. Overall, these results suggest that deletion of the MTS in *P. aeruginosa* RNase E modifies the physical properties and dynamics of RNase E-driven biomolecular condensates without abolishing their formation, thus maintaining a degree of RNA degradosome compartmentation.

Contrastingly, previous studies performed in *E. coli* showed that deletion of the MTS not only results in delocalisation of RNase E from the membrane but also completely abolishes foci formation (Strahl *et al*., 2015; Thappeta *et al*., 2025), highlighting the need of membrane anchoring for foci assembly or stability in this organism. The difference between *E. coli* and *P. aeruginosa* MTS deletion mutants likely arises from species-specific properties of the RNase E C-terminal scaffolding domain (Aït-Bara *et al*., 2015). In a previous study, we have described a REER-repeats region within *P. aeruginosa* RNase E C-terminal domain that is highly rich in arginine (R) and glutamate (E) residues (Geslain *et al*., 2025). This region, which is necessary for foci formation and binds RNA, is completely absent from the *E. coli* RNase E homolog (Aït-Bara *et al*., 2015). Interestingly, alphaproteobacterial RNase E homologs do not contain an MTS but drive the formation of biomolecular condensates owing to alternating blocks of negatively or positively charged residues, suggesting that the REER-repeats region may fulfil a similar function in *P. aeruginosa* and bypass the need of the MTS for maintenance of RNA degradosome compartmentation.

Consistent with the different effect of the RNase E MTS deletion on RNA degradosome organisation between *E. coli* and *P. aeruginosa*, distinct transcriptome profiles are observed in the two species. In *E. coli*, loss of the MTS results in widespread mRNA stabilization affecting more than 240 transcripts (Hadjeras *et al*., 2019; Moffitt *et al*., 2016), indicating a broad disruption of RNA turnover. In contrast, the *P. aeruginosa rne*ΔMTS mutant displays a relatively limited set of misregulated transcripts at steady state (n=93), suggesting that RNA degradosome compartmentation and its associated enzymatic activity are only partially affected. Transcripts encoding proteins with predicted transmembrane helices are enriched among the upregulated mRNAs in the *P. aeruginosa rne*ΔMTS mutant. A similar enrichment has been reported in *E. coli*, where deletion of the RNase E MTS preferentially stabilizes inner membrane-associated transcripts (Moffitt *et al*., 2016). In that model, co-translational insertion of nascent peptides localises mRNAs encoding inner membrane proteins near the membrane, in the vicinity of RNase E (Moffitt *et al*., 2016). Deletion of the RNase E MTS disrupts this spatial coupling and results in selective stabilisation of transcripts normally more rapidly decayed at the membrane. We hypothesise that a comparable mechanism may occur in *P. aeruginosa*. To confirm the subcellular localisation of membrane protein-encoding mRNAs affected in the *P. aeruginosa rne*ΔMTS, we performed smFISH. The putP transcript, which steady-state levels accumulate in the *rne*ΔMTS mutant, shows clear membrane enrichment, consistent with the proximity-based model in which membrane-associated transcripts are preferentially stabilised upon MTS deletion. In contrast, the pqsAB transcript does not display strong membrane localisation despite encoding an inner membrane protein, and its lack of enrichment correlates with reduced transcript levels in the mutant. Although additional transcripts will need to be analysed, these observations support the current model in which spatial coupling between mRNA localisation and RNA degradosome positioning contributes to a selective regulation of membrane-associated transcripts. The determinants of this selection are currently unclear.

Beyond an impact on specific transcript levels, our study demonstrates that altered RNA degradosome organisation leads to specific phenotypic impairments in *P. aeruginosa*. Deletion of the RNase E MTS results in a pronounced growth defect under hypersaline conditions induced by NaCl or KCl, whereas sucrose does not impair growth relative to the WT strain, suggesting a sensitivity to ionic stress rather than osmotic pressure per se. The mutant also exhibits mild cold sensitivity; a phenotype commonly associated with mutations in core RNA-binding proteins. Moreover, the modest but reproducible attenuation of virulence further implicates MTS-dependent RNA degradosome spatial organisation in host adaptation. These phenotypes may arise from disrupted processing of membrane-localized mRNAs in the MTS-deleted background. Alternatively, redistribution of RNase E within the cell may bring it into contact with RNA populations that are normally spatially segregated from the RNA degradosome, resulting in inappropriate RNA processing or decay and impaired stress adaptation. In *E. coli*, various small RNAs become strongly enriched at the cell poles during osmotic stress, suggesting that the partial polar relocalisation of RNase E observed in the *rne*ΔMTS mutant could perturb their expression levels (Kannaiah *et al*., 2019). Altogether, these findings establish a direct functional link between RNA degradosome spatial organisation and bacterial stress adaptation.

To determine whether the MTS-mediated control of RNA degradosome foci assembly, dynamics, and subcellular localisation depends primarily on amphipathicity or on specific sequence features within the MTS, we replaced the native MTS with a heterologous amphipathic helix. This approach allowed us to experimentally distinguish phenotypes caused by mere loss of amphipathicity from sequence-specific effects. Amphipathicity alone was sufficient to restore RNase E membrane localisation, re-establish RNA degradosome organisation into transient foci, and rescue stress adaptation. However, incomplete restoration of foci morphology and dynamics in the chimeric strain indicates that the native MTS sequence possesses intrinsic features which can modulate RNA degradosome behaviour beyond mere membrane targeting. Differences in primary sequence may influence the strength or stability of membrane association, thereby affecting spatial compartmentation and higher-order organisation of the RNA degradosome. Alternatively, the MTS may also participate in regulatory protein–protein interactions. In *E. coli*, the oscillating protein MinD transiently interacts with the RNase E MTS; its deletion results in frequent polar accumulation of RNase E and a significant impairment of RNase E dynamics (Kannaiah *et al*., 2024), echoing some of the alterations we observed in the *P. aeruginosa rne*ΔMTS mutant.

Overall, our study establishes the RNase E MTS as a multifunctional determinant of RNA degradosome organisation in *P. aeruginosa*. Beyond acting as a membrane anchor, the MTS modulates bacterial stress adaptation and virulence through its control of RNA degradosome subcellular distribution, dynamics, and mRNA processing and steady-state levels. Our findings demonstrate that amphipathic helices are not mere anchors and can encode regulatory information. More broadly, our study highlights subcellular organisation as a fundamental layer of bacterial gene regulation and illustrates how cellular architecture shapes adaptive gene expression.

## Acknowledgements

We would like to thank Prof. Christoph Engl for advice concerning the FISH protocol.

## Materials and Methods

### Bacterial strains and growth conditions

Bacterial strains used in this study are listed in Appendix Table S1. Unless otherwise specified, growth conditions were as described in Geslain et al. (2025).

### Plasmid construction

Details about primers used in this study and plasmid construction can be found in Appendix Tables S2 and S3, respectively.

### Construction of *P. aeruginosa* PAO1 mutant strains

Mutant strains were constructed identically as in Geslain et al. (2025).

### Western Blot

Western Blots were performed following the exact protocol described in Geslain et al. (2025).

### Microscopy imaging

For all microscopy experiments, bacteria were grown until OD_600nm_= 0.3-0.8 in 3 ml of NYB. When needed, membrane staining with FM4-64 was done by adding 0.3 μL of 10 mM FM4-64 (in DMSO) to 100 μL of culture in an Eppendorf tube. An incubation was done at 37°C for 10 minutes (no shaking) before proceeding with imaging. The cells were deposited on an agarose pad containing 1% UltraPure LMP agarose (Invitrogen) dissolved in 0.5X PBS. Phase contrast and widefield fluorescence microscopy images were acquired on a Zeiss Axio Imager 2 equipped with an objective alpha Plan-Apochromat 100x/1,46 Oil Ph3 M27 and a ZEISS Axiocam 305 mono CCD camera.

Structured Illumination Microscopy (SIM) fluorescence images were acquired with the ELYRA7 microscope (Zeiss) using excitation lasers of 488 nm (for msfGFP), 561 nm (for mCherry or FM4-64) or 405 nm (for mTagBFP2). Laser intensities of 6-8% and exposure times of 60-80 ms were used for imaging of fluorescently-tagged proteins, while 0.5-1% laser was sufficient to visualise the FM4-64 membrane dye.

For time-lapse acquisition, an interval of 5000 ms between each frame and a total of 25 cycles were chosen. The software definite focus function was used to prevent loss of focus over time. Reconstruction was done using the Processing tab of the Zen Blue software (SIM^2^ algorithm) using the parameters shown in Appendix Fig S4.

### Image processing

All image treatments were performed using Fiji (Schindelin *et al*, 2012). Representative microscopy images and images to be analysed were opened in Fiji and cropped when needed to keep only cells showing good in-focus signal. Background noise was measured in an area devoid of cells and the mean background signal value was subtracted from the entire image by using the Process -> Math -> Subtract function. When showing merged representative images, merging of the separated phase-contrast and fluorescence channels was done using the Image -> Color -> Merge Channels function in Fiji. When a shift between the phase contrast channel and the fluorescence channel was observed (e.g. several foci or cell background signal being outside of phase contrast cells all in the same direction), this was manually corrected for the entire image using the Image -> Transform -> Translate function in Fiji. Display was adjusted using the Reset function in the Adjust Brightness/Contrast menu for most images. For FISH images, the display range was manually set to be identical between images of the WT and the *rne*ΔMTS strain (and also with the negative control when possible).

### Growth assays on solid media

For solid medium growth assays, square Petri dishes (12×12cm) were filled with 50 mL of warm nutrient agar (NA), prepared with 40 g Oxoid blood agar base and 5 g Difco yeast agar per litre. For testing growth with 0,75M NaCl or KCl, 17.54 g of NaCl or 22.36 g of KCl were added to 400 mL of freshly prepared NA medium and autoclaved before pouring into square Petri dishes. For testing growth with 0.75M sucrose, a 2X NA concentrate was prepared and mixed with an equal volume of 1.5 M sucrose solution when still hot before pouring into a square Petri dish. The Petri dishes were left to dry on the bench for 24-36h before use.

For the spotting assay, fresh cultures (OD_600nm_= 0.4-0.8) were adjusted to OD_600nm_=1 by centrifugation and serially diluted in a 96-well plate using a multipipette. 5 μL of each dilution series were then spotted using a multipipette onto pre-dried NA plates. Plates were incubated upside down at 37°C or 16°C for approximately 24h or 48h, respectively.

### *Galleria mellonella* killing assays

*G. mellonella* infections were conducted following the exact protocol described in Geslain et al. (2025).

### RNA-seq sample collection

The RNA-seq sample collection was done following the exact protocol described in Geslain et al. 2025. Of note, the *rne*ΔMTS strain was grown and harvested in parallel to the WT strain whose data is already published in Geslain et al. (2025). The analysis was thus done comparing the *rne*ΔMTS dataset to the RNA-seq dataset of PAO1 WT from Geslain et al. (2025).

### RNA sequencing analysis

The sequencing data was processed and mapped exactly as described in Geslain et al. (2025).

### RT-qPCR

RT-qPCR verification of relative transcript abundance in the WT or *rne*ΔMTS RNA samples was done following the exact protocol described in Geslain et al. 2025. The genes rpoD, putP, asrA, pqsA, mexX and kdpA were targeted using the primer pairs q65-q66, q116-q117, q120-q121, q112-q113, q118-q119, and q114-q115, respectively. Primer sequences are disclosed in Appendix Table S2.

### GC-EMOTE library construction

The GC-EMOTE experiment was performed starting from the same RNA samples used for RNA sequencing. The protocol published by Redder (2018) as used as a template and optimised for GC-rich organisms like *P. aeruginosa* as follows (Redder, 2018). The primers and oligos used for the GC-EMOTE experiment are listed in Appendix Table S4.

To begin with, TURBO DNase-treated gDNA-free total RNA samples were depleted for rRNA using the riboPOOL kit, following the manufacturer’s instructions and using 5 μg of total RNA as starting material. After rRNA depletion, the RNA samples were purified using the ethanol precipitation method, replacing the linear acrylamide recommended in the kit by GlycoBlue (1 μL per sample). After all the washing steps, drying of the RNA pellet was done for 5 minutes under a clean sterile flowhood and the pellet was resuspended in 10 μL of nuclease-free 0.1X TE buffer. For ligation of the BioVm1 RNA oligo, a ligation mixture was prepared (per sample) as follows: 2 μL 10X T4 RNA ligase, 2 μL of 10 mM ATP, 0.8 μL RNasin Plus (Promega), 4.2 μL PEG8000 50% (NEB), and 1 μL T4 RNA ligase (NEB). A mixture of 9 μL of rRNA-depleted RNA sample plus 1 μL of 100 μM BioVm1 was prepared, heated to 80°C for 2 minutes and flash-cooled on ice. The ligation mixture was then added to the sample and incubated for > 16h at 16°C. Ethanol precipitation was then done as described in Redder (2018) with minor changes: two volumes of ethanol were added instead of three and washing was done with 400 μL of cold 70% ethanol. Final resuspension of the pellet, after 5 minutes drying under the sterile flowhood, was done in 10 μL nuclease-free 0.1X TE buffer.

For the reverse-transcription reaction, the ImProm-II reverse transcriptase was used (Promega). The MasterMix was prepared according to the manufacturer’s instructions and 1 μL of 20 μM DROCC primer was mixed with 9 μL ligated RNA and heated for 2 min at 80°C before allowing a slow cooling to room temperature (5 min with tubes lying on top of turned off heat block and 5 min in an Eppendorf rack). 10 μL of Reverse Transcription mix were added to each sample of RNA mixed with DROCC primer. Tubes were left at room temperature for 10 min then incubated at 42°C for 60 min before heat-inactivating the reverse transcriptase at 70°C for 15 min. The cDNA was purified with AMPure XP beads (Beckman Coulter) using a 1.5X ratio of beads to sample and following the manufacturer’s instructions. Elution was done in 40 μL of nuclease-free 0.1X TE buffer.

For PCR amplification and tagging, a Mastermix was prepared (per sample) as follows: 7.5 μL of mQH_2_O, 10 μL of 5M betaine, 1.5 μL of DMSO, 10 μL of 5X Q5 reaction buffer, 4 μL of dNTPs, 2.5 μL of 10 μM primer E2, and 0.5 μL of Q5 High Fidelity DNA polymerase. After splitting 36 μL of MasterMix into different PCR tubes, 10 μL of purified cDNA sample and 2.5 μL of 100 nM primer pVmA, pVmB, pVmC, pVmD, pVmE, pVmF, pVmG, pVmH, pVmI, pVmJ, pVmK, or pVmL (one primer for each cDNA sample) were added per tube to allow the generation of a unique tag for each sample. A first short touchdown PCR was run with the following programme: 98°C for 2 minutes; 5 cycles of 98°C for 10s, 67°C (1x) / 66°C (1x) / 65°C (1x) / 64°C (1x) / 63°C (1x) for 30s, 72°C for 45s. At the end of the programme, PCR tubes were quickly placed on ice, supplemented with 2.5 μL of 10 μM primer E1, and run with a second touchdown PCR programme: 98°C for 1min 50s; 25 cycles of 98°C for 10s, 67°C (1x) / 66°C (1x) / 65°C (1x) / 64°C (1x) / 63°C (21x) for 30s, 72°C for 1 minute; 72°C for 5 min. At the end of the run, the PCR products were purified with AMPure XP beads (Beckman Coulter) using a 0.8X ratio of beads to sample after pooling the barcoded PCR products per series of replicates. At the end of the purification, the dsDNA was eluted in mQH_2_0 and stored at −20°C. Sequencing was done on the Novaseq X Plus Series (PE150) platform at Novogene (120 G raw data per sample).

### GC-EMOTE sequencing data processing and mapping

Raw sequencing data were demultiplexed by barcode using custom Bash scripts, trimming the four nucleotide barcodes from the 5’ end as they were identified, along with the single random nucleotide upstream. Adapter trimming was then performed on the remaining downstream read using cutadapt v4.9 (Martin, 2011). A 19-nucleotide overlap with zero error rate was required, and the 5′ fixed tag sequence CTTCACCACACCCTATCAC was removed from reads. Only trimmed reads were retained for downstream analysis. Unique molecular identifiers (UMIs) were extracted from the 5′ ends of reads using UMI-tools v1.1.6 with a regular expression capturing the 17-nucleotide UMI (excluding guanine) immediately upstream of the fixed ACATA motif (Smith *et al*, 2017). Reads were further length-trimmed to 25 nucleotides using cutadapt for consistent downstream alignment.

Trimmed reads were aligned to the *Pseudomonas aeruginosa* PAO1 genome assembly ASM676v1 using bowtie2 v2.5.4 in end-to-end, very-sensitive mode (Langmead & Salzberg, 2012). SAM alignments were converted to BAM format, sorted, and indexed with samtools v1.20 (Li *et al*, 2009). PCR duplicates were removed using UMI-tools with the directional method, leveraging UMI sequences to identify and collapse duplicate reads. Five prime end depths were calculated genome-wide, and quantification relative to transcript features used ASM676v1 gtf version 2.2 (NCBI accession: GCF_000006765.1) for locations.

### Single molecule fluorescence in situ hybridisation (smFISH)

TAMRA-labelled ssDNA probes for FISH were synthesised by the Stellaris RNA FISH probes service (Biosearch Technologies). A set of 47 to 48 probes was used to target putP, asrA, or pqsAB mRNAs (Appendix Table S5). Of note, pqsAB was chosen instead of pqsA alone due to the difficulty of getting a sufficient number of specific probes binding in pqsA only. The Stellaris FISH probe designer was used as a base generator of potential FISH probes (18-, 19- or 20-mers), after which all the candidate probes were aligned against the *P. aeruginosa* genome (NC_002516) using Snapgene. Probes with poor specificity (approx. > 20-30 potential binding sites) or with unspecific sense strand targets having a predicted Tm of > 50°C were typically excluded from the dataset. A minimal spacing of two nucleotides between each probe was chosen, with a preferential selection of probes allowing a spacing or three or more. Finally, the probe length was manually adjusted to achieve similar Tm values for most probes based on PerlPrimer calculations (Marshall, 2004).

The FISH protocol from Skinner et al. (2013) was used as a base protocol. All reagents were as described in the published protocol unless otherwise specified (Skinner *et al*., 2013). A few changes were made to the protocol, as described hereafter.

Fresh cultures were started in 10 mL of NYB broth (100 mL Erlenmeyer) for asrA / pqsAB, or in 15 mL of NYB broth (250 mL Erlenmeyer) for putP. 1 mL of overnight culture was centrifuged, resuspended in fresh NYB, and diluted 1000-fold for inoculation of the fresh cultures. The cultures were cross-linked when OD_600nm_ was ≈0.4 for FISH against putP or ≈1 for FISH against pqsAB / asrA. A 5X fixing solution containing a 1:1 mix of 37% formaldehyde (Sigma) and 1X RNase-free PBS (ThermoFisher Scientific) was directly added to the cultures for cross-linking. The Erlenmeyers were sealed with parafilm and incubated for 15 minutes at 37°C with shaking, followed by a 30 minutes static incubation on ice. A volume of culture corresponding to *V*(ml) = 4/OD_600nm_x1.2 was then transferred into 50 mL Falcon tubes and centrifuged at 800g with slow braking, 4°C, for 20-25 minutes. The subsequent washing and permeabilization steps were performed as described in Skinner et al. (2013), increasing the centrifugation times from 3.5 min to 6 min. The hybridisation solution was prepared as described in Skinner et al. (2013), with minor changes: the concentration of ribonucleoside-vanadyl complex was increased from 2 mM to 10 mM and *S. cerevisiae* tRNA (Sigma-Aldrich) was used instead of *E. coli* tRNA. FISH probes were briefly denatured by heating at 80°C for 3 minutes and placing them on ice before adding them to the hybridisation mix. After resuspension of the permeabilised cells into the hybridisation mix, the suspension was heated at ≈ 67°C for 5 minutes in a heat-block, plus an additional 10 minutes after turning off the heat block. A slow 5-minutes cool-down was then done by leaving the tubes lying on top of the heat block before placing them in an Eppendorf shaker at 30°C overnight (450 rpm shaking). Of note, FISH probes and suspensions containing FISH probes were shielded from light at all times.

The next day, washing steps were performed mostly as described in the published protocol, reducing the incubation times to 20 minutes and performing an additional final 10-minutes wash in 2x SSC before putting ≈1-2 μL of suspension onto a pad made with 0.5X RNase-free PBS (ThermoFisher Scientific) and 1% TopVision agarose (ThermoFisher Scientific). Visualisation was done using epifluorescence microscopy (800ms exposure time for the TAMRA channel).

### RNase E foci or smFISH foci quantification / subcellular localisation analysis

All image treatments and analysis were performed using Fiji (Schindelin *et al*., 2012). Images to be analysed were first opened in Fiji and processed as described in the Image Processing section.

Detection of local maxima and their assignment to the detected bacterial particle was done with MicrobeJ (Ducret *et al*, 2016). Subsequent quantification of foci counts per cell, foci intensities (or other characteristics) and generation of average local maxima subcellular localisation were also generated with the MicrobeJ plugin (Ducret *et al*., 2016). Parameters used for bacteria and maxima detection of images showing either msfGFP-tagged RNase E variants or mRNAs visualised by FISH are shown in Appendix Fig. S5. Manual segmentation of cells was done in some cases to avoid excluding many cells from the analysis, always manually verifying that the foci count has not been affected for segmented cells. Erroneous automatic detection of cells was manually corrected by removing those elements from the results table. The results were then exported as a .csv file. Data obtained for several individual images across three biological replicates was pooled by pasting all the values into GraphPad Prism. Graphs of foci intensity and other foci morphology measurements were then plotted using GraphPad Prism. Graphs showing the average foci subcellular localisation (RNase E foci or FISH foci) were generated directly with MicrobeJ. The “XYCellDensity” graphing method was selected, and the density bandwith settings used were x = 0.05 and y = 0.25 for all graphs generated. The default look-up table (“fire”) was chosen for all graphs. Images were saved as .svg files and colours were inverted in Adobe Illustrator with the Edit Colors -> Invert Colors function.

---

**Figure EV1:**
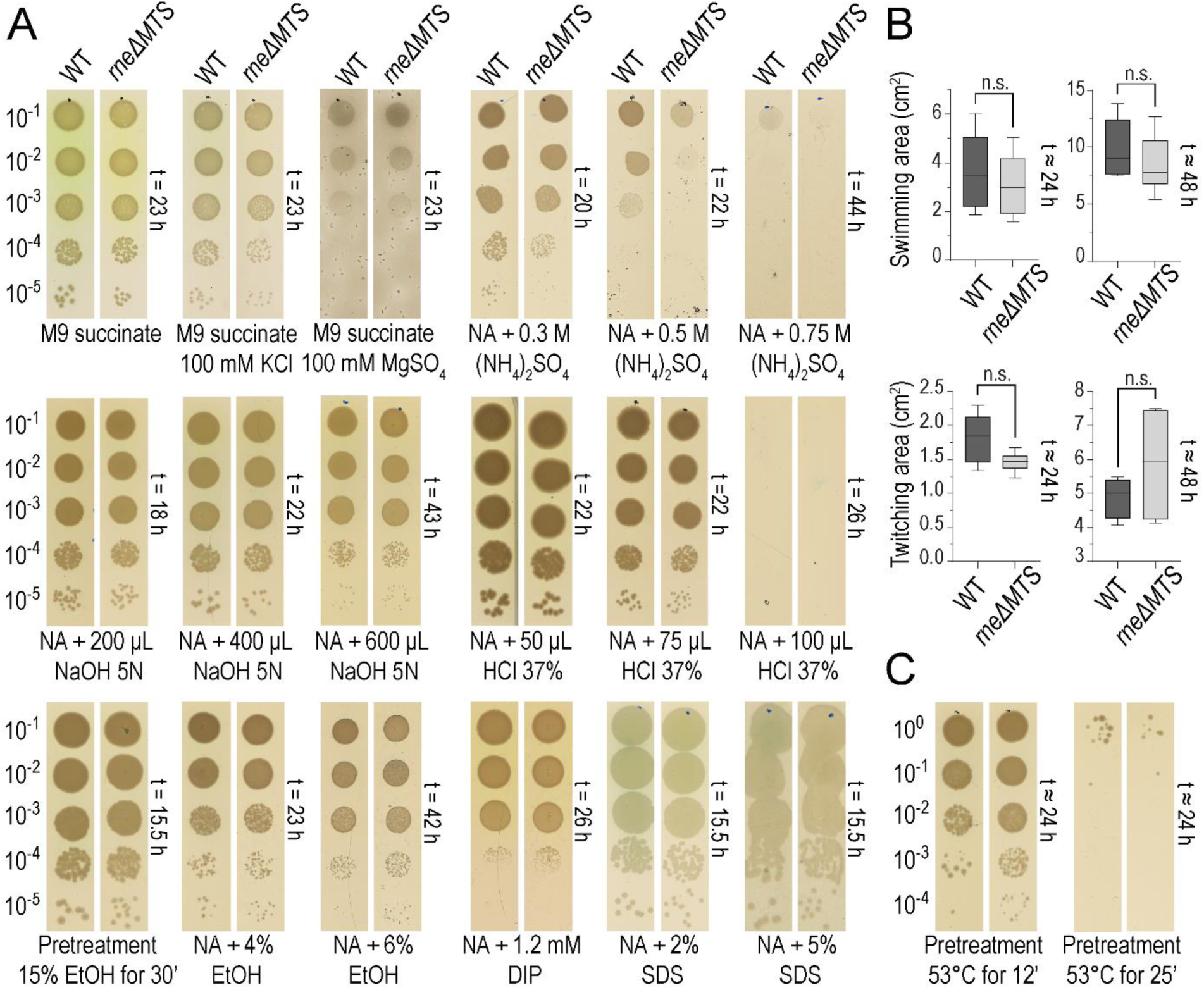
Extensive phenotypic screening of the *rne*ΔMTS strain reveals a specific sensitivity to high salinity or acidic stress but no broad fitness cost. (A) Spot assay screening of *P. aeruginosa* WT or *rne*ΔMTS growth on various solid media as indicated below each image. Time of imaging, recorded as hours post-inoculation, is indicated for each plate. Images shown are representative of one to two biological replicates. (B) Swimming and twitching motility assays quantified after approximately 24h or 48h post-imoculation. Two to three independent biological replicates were performed and pooled for plotting. Statistical significance was assessed with a two-tailed unpaired t-test with Welch’s correction. n.s.: not significant. (C) Spot assay on standard NA plates with the *P. aeruginosa* WT (left) or *rne*ΔMTS (right) strains after incubation of 100 μL of cultures in Eppendorf tubes in a 53°C water bath for 12 or 25 minutes.

**Figure EV2:**
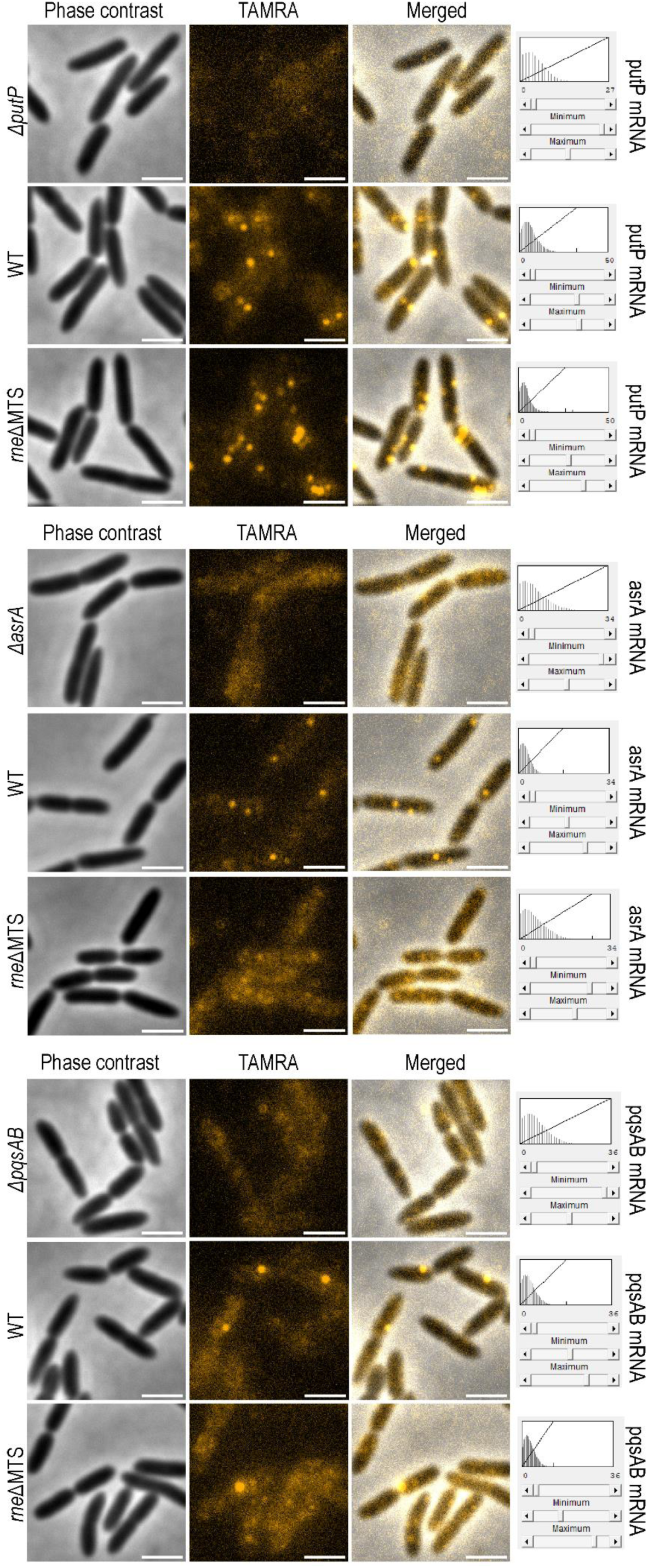
Visualisation of FISH signal by epifluorescence microscopy. The phase contrast channel, fluorescence channel, or merged images, are shown separately. For each acquisition, the strain background is indicated on the left and the mRNA visualised is indicated on the right. Scale is 2 μm. The display range, adjusted in Fiji for each fluorescence image, is shown for reference.

