## Supplemental information for "Intrinsic features of the RNase E membrane targeting sequence specify RNA degradosome organisation and activity"

### Appendix

Appendix Table S4 ........ 4-5
Appendix Table S5 ........ 5-8

**Appendix Figures** ........ **9-13**

**Appendix Table S1:** Bacterial strains used in this study.

| Strains | Genotypes / description | Source |
|---|---|---|
| *E. coli* | | |
| DH5 | recA1 endA1 hsdR17 supE44 thi-1 gyrA96 relA1 Δ(lacZYA-argF)U169 [Φ80dlacZM15]F- NaI | |
| *P. aeruginosa* | | |
| PAO1 | Wild-type | |
| PAO1 *rne::2xStrep* | Chromosomal insertion of a Twin Strep-tag immediately upstream of the stop codon in *rne* | Geslain et al., 2025 |
| PAO1 *rne::msfGFP* | Chromosomal insertion of msfGFP with a flexible linker immediately upstream of the stop codon in *rne* | Geslain et al., 2025 |
| PAO1 *rneΔMTS* | In-frame deletion of *rne* sequence coding for residues 552-573 (MTS and a few downstream residues) in the PAO1 WT strain | This study |
| PAO1 *rneΔMTS::2xStrep* | In-frame deletion of *rne* sequence coding for residues 552-573 (MTS and a few downstream residues) in the PAO1 rne::2xStrep strain | This study |
| PAO1 *rneΔMTS::msfGFP* | In-frame deletion of *rne* sequence coding for residues 552-573 (MTS and a few downstream residues) in the PAO1 rne::msfGFP strain | This study |
| PAO1 *rneMinDα* | In-frame insertion of the MinD amphipathic helix (MinDα) in the PAO1*rneΔMTS* at the native MTS position | This study |
| PAO1 *rneMinDα::2xStrep* | In-frame insertion of the MinD amphipathic helix (MinDα) in the PAO1*rneΔMTS::2xStrep* at the native MTS position | This study |
| PAO1 *rneMinDα::msfGFP* | In-frame insertion of the MinD amphipathic helix (MinDα) in the PAO1*rneΔMTS::msfGFP* at the native MTS position | This study |
| PAO1 *rhl::msfGFP/ rne::mCherry /pnp::mTagBFP2* | Chromosomal insertion of *mTagBFP2* with a flexible linker immediately upstream of the pnp stop codon in the strain PAO1 rhl::msfGFP/ rne::mCherry (from Geslain et al. 2025). | This study; Geslain et al. 2025 |
| PAO1 *rhl::msfGFP/ rneΔMTS::mCherry /pnp::mTagBFP2* | Deletion of the *rne* sequence coding for residues 552-573 (MTS and a few neighboring residues) in the PAO1 *rhl::msfGFP/ rne::mCherry* / *pnp::mTagBFP2* strain | This study |
| PAO1 *rhl::msfGFP/ rneMinDα::mCherry /pnp::mTagBFP2* | In-frame insertion of the MinD amphipathic helix (MinDα) in the PAO1 *rhl::msfGFP/ rneΔMTS::mCherry* / *pnp::mTagBFP2* strain at the native MTS position | This study |
| PAO1 *ΔputP* | Deletion of the entire putP CDS and partial deletion of intergenic regions or UTRs (genomic positions 855224-856801) | This study |
| PAO1 *ΔasrA* | In-frame deletion of most of the asrA CDS ( genomic positions 845820- 848183) | This study |
| PAO1 *ΔpqsAB* | In-frame deletion of most of the pqsA and pqsB CDS (genomic positions 1078468 - 1080785) | This study |

**Appendix Table S2:** Primers used in this study.

| Primer code | Primer sequence (5' -> 3') |
|---|---|
| p399 | CAATATAGGAGCATAGAATTCTCCAGGTCGGTCGCATCTC |
| p400 | CTTGCTGGTTTCAGCAGGCTTCGGCTCGGGCATCGGCTT |
| p401 | AAGCCGATGCCCGAGCCGAAGCCTGCTGAAACCAGCAAG |
| p402 | GAGAATGGCAAAAGCTTTCGCCATCGGTATCGTCCTG |
| p801 | GTCGACTCTAGAGGATCCGACATCAAGATCAACGGTATCACCGAAG |
| p802 | GAACCACCACCCACGCCGGAAGCTTCGG |
| p803 | CGTGGGTGGTGGTTCCGTGTCTAAGGGCGA |
| p804 | ACGACTCGACTTAATTAAGCTTGTGCCCCAGTTTGCT |
| p805 | GCTTAATTAAGTCGAGTCGTGATCGATGAAAACAG |
| p806 | GAAATTAATTAAGGTACCGAATTCGGCTTTCACGCCCTTGATGC |
| p971 | CAGGTCGACTCTAGAGGATCCCAGGTCGGTCGCATCT |
| p976 | TTAATTAAGGTACCGAATTCTCGCCATCGGTATCGTCCTG |
| p1007 | TCCTCCGAACAAGCGTTTGAGGAAGCCTTTCTTCGGCTCGGGCATCGGCTTG |
| p1008 | AAGAAAGGCTTCCTCAAACGCTTGTTCGGAGGAAAGCCTGCTGAAACCAGCAAGC |
| p1037 | CAGGTCGACTCTAGAGGATCCCGACGAGCTGAAGAAAGAGAACTTC |
| p1038 | CGCCCGCAGGCGGTAGCGGGCCCTTGGTTG |
| p1039 | AGGGCCCGCTACCGCCTGCGGGCGGGTAAAG |
| p1040 | GGAAATTAATTAAGGTACCGAATTCCTCTGCCTGTTCGACGAATGC |
| p1074 | CAGGTCGACTCTAGAGGATCCAGGTTCCGCACGATATGC |
| p1075 | ACCTTGGCCACGTCGCTCATGAGGCACCT |
| p1076 | CCTCATGAGCGACGTGGCCAAGGTGCTGT |
| p1077 | GGAAATTAATTAAGGTACCGAATTCCGCGCAGGAAAAGCTTTC |
| p1078 | CAGGTCGACTCTAGAGGATCCAGGTGTCCTCTTCGGCAG |
| p1079 | TCGAAGCTGATCAGGGACATGACAGAACGTTCCC |
| p1080 | TTCTGTCATGTCCCTGATCAGCTTCGATCCGTTC |
| p1081 | GGAAATTAATTAAGGTACCGAATTCGCACACCACCAGCACTTTCTC |
| q65 | CGTCGCGGCTACAAGTTCTCCA |
| q66 | GGCTCGCGACCCATTTCCTG |
| q112 | GATCCCGATACCGCCGTTTATCACTATC |
| q113 | GCGGGACTTGGGATTGATCACG |
| q114 | GCTTCAAGGTGATGGAAGGCCAAC |
| q115 | GCCAGGGTGTAGGTCTTCCAGTC |
| q116 | CTCCACCAACAACTTCTCCGACTACATC |
| q117 | TTGCCGTTGTGCTCGGTCTG |
| q118 | GCTGGGCTGCGAAGAAGCA |
| q119 | CGTCCTGGCCTTCCTCGTACA |
| q120 | GGATAGTCCACCTCCACCATGAACG |
| q121 | TTCATGGACAATCCTCCGGAAGACC |

**Appendix Table S3:** Plasmids used in this study.

| Name | Used for: | Insert(s) amplified with: | Template used in PCR |
|---|---|---|---|
| pME3087_RNase E$^{\Delta MTS}$ | Construction of PAO1 *rneΔMTS; rneΔMTS::2xStrep*; *rneΔMTS::msfGFP*; *rhl::msfGFP/ rneΔMTS::mCherry /pnp::mTagBFP2* | p399-p400; p401-p402 | PAO1gDNA |
| pEXG2_RNase E$^{MinD\alpha}$ | Construction of PAO1 *rneMinDα; rneMinDα::2xStrep*; *rneMinDα::msfGFP; rhl::msfGFP/ rneMinDα::mCherry /pnp::mTagBFP2* | p971-p1007; p1008-p976 | PAO1gDNA |
| pEXG2_PNPase-mTagBFP2 | Construction of PAO1 *rhl::msfGFP/ rne::mCherry /pnp::mTagBFP2* | p801-p802; p803-p804; p805-p806 | *pnp*-containing plasmid; mTagBFP2-containing plasmid |
| pEXG2_ΔPutP | Construction of PAO1 *ΔputP* | p1037-p1038; p1039-p1040 | PAO1gDNA |
| pEXG2_ΔAsrA | Construction of PAO1 *ΔasrA* | p1074-p1075; p1076-p1077 | PAO1gDNA |
| pEXG2_ΔPqsAΔPqsB | Construction of PAO1 *ΔpqsAB* | p1078-p1079; p1080-p1081 | PAO1gDNA |

**Appendix Table S4:** Oligos and primers used for preparation of the GC-EMOTE library.

| Name | Sequence (5' -> 3') | Oligo type | Ordered from | Purification |
|---|---|---|---|---|
| BioVm1 | Biotin-CUUCACCACACCCUAUCACHHHHHHHHHHHHHHHHACAUA | RNA | IDT | Standard Desalting |
| DROCC_i7 | CTGGAGTTCAGACGTGTGCTCTTCCGATCTNNNNNNNNNNCC | ssDNA | IDT | RNase-Free HPLC |
| pVmA | CTCTTTCCCTACACGACGCTCTTCCGATCTNTACACTTCACCACACCCTATCAC | ssDNA | Microsynth | IEX-HPLC |
| pVmB | CTCTTTCCCTACACGACGCTCTTCCGATCTNGTATCTTCACCACACCCTATCAC | ssDNA | Microsynth | IEX-HPLC |
| pVmC | CTCTTTCCCTACACGACGCTCTTCCGATCTNCGTCCTTCACCACACCCTATCAC | ssDNA | Microsynth | IEX-HPLC |

| | | | | |
|---|---|---|---|---|
| pVmD | CTCTTTCCCTACACGACGCTCTTCCGATCTNAAGTCTTCACCACACCCTATCAC | ssDNA | Microsynth | IEX-HPLC |
| pVmE | CTCTTTCCCTACACGACGCTCTTCCGATCTNACACCTTCACCACACCCTATCAC | ssDNA | Microsynth | IEX-HPLC |
| pVmF | CTCTTTCCCTACACGACGCTCTTCCGATCTNGGTACTTCACCACACCCTATCAC | ssDNA | Microsynth | IEX-HPLC |
| pVmG | CTCTTTCCCTACACGACGCTCTTCCGATCTNTCGGCTTCACCACACCCTATCAC | ssDNA | Microsynth | IEX-HPLC |
| pVmH | CTCTTTCCCTACACGACGCTCTTCCGATCTNCAAGCTTCACCACACCCTATCAC | ssDNA | Microsynth | IEX-HPLC |
| pVmI | CTCTTTCCCTACACGACGCTCTTCCGATCTNTTGACTTCACCACACCCTATCAC | ssDNA | Microsynth | IEX-HPLC |
| pVmJ | CTCTTTCCCTACACGACGCTCTTCCGATCTNGCTGCTTCACCACACCCTATCAC | ssDNA | Microsynth | IEX-HPLC |
| pVmK | CTCTTTCCCTACACGACGCTCTTCCGATCTNCCGACTTCACCACACCCTATCAC | ssDNA | Microsynth | IEX-HPLC |
| pVmL | CTCTTTCCCTACACGACGCTCTTCCGATCTNCTCGCTTCACCACACCCTATCAC | ssDNA | Microsynth | IEX-HPLC |
| E1 | AATGATACGGCGACCACCGAGATCTACACTCTTTCCCTACACGACG | ssDNA | Microsynth | IEX-HPLC |
| E2 | CAAGCAGAAGACGGCATACGAGATCGTGATGTGACTGGAGTTCAGACGTGTGC | ssDNA | Microsynth | IEX-HPLC |

**Appendix Table S5:** List of TAMRA-labelled FISH probes against putP, pqsAB, or asrA. Tm values were calculated using PerlPrimer (Marshall, 2004).

| FISH probe name | FISH probe sequence | Length | GC content | Predicted Tm |
|---|---|---|---|---|
| putP_probe_1 | GCGCCGTGACGAAGCT | 16-mer | 68% | 63.44°C |
| putP_probe_2 | TGGCCGAAGTAGCCCAG | 17-mer | 64% | 61.55°C |
| putP_probe_3 | CAGCCGCTCATATCGGAG | 18-mer | 61% | 60.18°C |
| putP_probe_4 | CGCGGTCTCGTAGGAGAG | 18-mer | 66% | 61.72°C |
| putP_probe_5 | AGGTCATGGAGATGCGGC | 18-mer | 61% | 62.34°C |
| putP_probe_6 | GATGCCGAAGAAGCCCAC | 18-mer | 61% | 61.28°C |
| putP_probe_7 | GGATGCGCCTGGAAGTAG | 18-mer | 61% | 60.34°C |
| putP_probe_8 | GAGGATGGCGGACAGCAA | 18-mer | 61% | 62.57°C |
| putP_probe_9 | CGAGGAGCAGACCAGCAG | 18-mer | 66% | 62.60°C |
| putP_probe_10 | CGGTTTTCCGGGTTGGAG | 18-mer | 61% | 61.13°C |
| putP_probe_11 | CAGGCATAGGACACCAGG | 18-mer | 61% | 59.54°C |
| putP_probe_12 | CGTTGTGCTCGGTCTGCA | 18-mer | 61% | 63.42°C |
| putP_probe_13 | GGTGAAGTAGTCGGGCAG | 18-mer | 61% | 59.87°C |
| putP_probe_14 | GGCTGTTGTCCTCGAAGC | 18-mer | 61% | 61.19°C |

| | | | | |
|---|---|---|---|---|
| putP_probe_15 | GGTGTCGGTCCAGCTCAC | 18-mer | 66% | 62.84°C |
| putP_probe_16 | CGAGCATCACGATCACCG | 18-mer | 61% | 61.10°C |
| putP_probe_17 | CTCGGGGTTCTCGCTGAC | 18-mer | 66% | 62.59°C |
| putP_probe_18 | CAGCCAGATGGCGATCAC | 18-mer | 61% | 61.04°C |
| putP_probe_19 | ACTCCTGCTCCGCATCGT | 18-mer | 61% | 63.92°C |
| putP_probe_20 | TCCACAGGATCACCGTCG | 18-mer | 61% | 61.53°C |
| putP_probe_21 | GGTCCAGCCCAACAGGTT | 18-mer | 61% | 62.38°C |
| putP_probe_22 | TTGCGTGTGCAAGCAACG | 18-mer | 55% | 63.00°C |
| putP_probe_23 | GATCCAGCTTTCGGACAGG | 19-mer | 57% | 60.62°C |
| putP_probe_24 | TGTTGGTGGAGCGATATGC | 19-mer | 52% | 60.62°C |
| putP_probe_25 | ACCGAGGATGTAGTCGGAG | 19-mer | 57% | 60.62°C |
| putP_probe_26 | GGAAGCGCAGTAGATGGTG | 19-mer | 57% | 60.99°C |
| putP_probe_27 | CACCGAGGCAGAGGATCAT | 19-mer | 57% | 61.62°C |
| putP_probe_28 | CGATCCACGGGTTGAACAG | 19-mer | 57% | 61.20°C |
| putP_probe_29 | AGCTCAGGGTGCTCATGAC | 19-mer | 57% | 62.15°C |
| putP_probe_30 | ACAGCAGGGAGAACAGCAC | 19-mer | 57% | 62.67°C |
| putP_probe_31 | GGGGATGATCTCGTACAGG | 19-mer | 57% | 59.11°C |
| putP_probe_32 | CGAAGGTGCTCTCGAACAG | 19-mer | 57% | 60.90°C |
| putP_probe_33 | GAACACCAGGATCACCAGG | 19-mer | 57% | 60.17°C |
| putP_probe_34 | CCCAGACCAGTTCCAGCTG | 19-mer | 63% | 62.74°C |
| putP_probe_35 | AGGCTGAAGACGACGATGG | 19-mer | 57% | 62.19°C |
| putP_probe_36 | TCGATCAGCGCTGGGTTAC | 19-mer | 57% | 62.55°C |
| putP_probe_37 | CCTTGTAGAAGTCCTCGGTG | 20-mer | 55% | 59.94°C |
| putP_probe_38 | GTGGGCGTATTGACACTCAT | 20-mer | 50% | 60.59°C |
| putP_probe_39 | GCGATGTAGATCACGAAGGT | 20-mer | 50% | 60.09°C |
| putP_probe_40 | CGAACAGCCAGTTGAGGTAG | 20-mer | 55% | 60.87°C |
| putP_probe_41 | CGATGAAGGTGTAGGCGATG | 20-mer | 55% | 61.07°C |
| putP_probe_42 | ATCAGCGCGAAGATCATCAG | 20-mer | 50% | 60.79°C |
| putP_probe_43 | CTTGAGCATGTCGAAGCTGG | 20-mer | 55% | 61.84°C |
| putP_probe_44 | GAGATCACGCCGATGAAGGA | 20-mer | 55% | 62.22°C |
| putP_probe_45 | GATCGATTTCACCGAGTCGG | 20-mer | 55% | 61.06°C |
| putP_probe_46 | CTTGAGCATGCTCGTCGAAG | 20-mer | 55% | 61.89°C |
| putP_probe_47 | GGACTCCCCTATTGCTTGTTT | 21-mer | 47% | 60.50°C |
| asrA_probe_1 | TTCGGGATTAACGTCCTGGT | 20-mer | 50% | 61.40°C |
| asrA_probe_2 | GGGAAGAACGGGCGGTTG | 18-mer | 66% | 63.85°C |
| asrA_probe_3 | CACCAGTTCGAGGGTTTCC | 19-mer | 57% | 60.77°C |
| asrA_probe_4 | AGCTCCTTGATCGCGTTGA | 19-mer | 52% | 62.12°C |
| asrA_probe_5 | GAAGTCGGTGAGTGGCGA | 18-mer | 61% | 62.14°C |
| asrA_probe_6 | CGCTCTCGAGGATCGACA | 18-mer | 61% | 61.35°C |
| asrA_probe_7 | TGATGCTTGTCGAGGACCTT | 20-mer | 50% | 61.69°C |
| asrA_probe_8 | ATGCGATCCTTGATGTCGTC | 20-mer | 50% | 60.73°C |
| asrA_probe_9 | GCCTTTTCCAACTGCTTGG | 19-mer | 52% | 60.10°C |
| asrA_probe_10 | AGGGTTCTTCGTTGACGATC | 20-mer | 50% | 60.23°C |

| | | | | |
|---|---|---|---|---|
| asrA_probe_11 | CGAATGGTGGTCGGTCTTG | 19-mer | 57% | 61.20°C |
| asrA_probe_12 | CAGGCTGTTGACGTCGAAG | 19-mer | 57% | 61.48°C |
| asrA_probe_13 | GTCGTTGGGACTGAAGCG | 18-mer | 61% | 61.18°C |
| asrA_probe_14 | CTCTTCGTCGATACGCTTGC | 20-mer | 55% | 61.67°C |
| asrA_probe_15 | GATGGATTTGCCGATGCTG | 19-mer | 52% | 59.80°C |
| asrA_probe_16 | GATCGAGTCGAGGGTGTTG | 19-mer | 57% | 60.32°C |
| asrA_probe_17 | GCAGTTTGACCACCGACTTG | 20-mer | 55% | 62.26°C |
| asrA_probe_18 | TGAGCTTGAAGCCACGGTT | 19-mer | 52% | 62.60°C |
| asrA_probe_19 | GGATTCCTTCATCACGTCGC | 20-mer | 55% | 61.63°C |
| asrA_probe_20 | GCCATACTTCTTCAGGTGCGA | 21-mer | 52% | 62.94°C |
| asrA_probe_21 | CACGTGCAGGTGGACGAA | 18-mer | 61% | 63.06°C |
| asrA_probe_22 | GGATCACGTAGAGGGTGGTC | 20-mer | 60% | 61.88°C |
| asrA_probe_23 | CGGAGGATTGTCCATGAAGAA | 21-mer | 47% | 60.29°C |
| asrA_probe_24 | ATCGGGGTCTTCGGATAGTC | 20-mer | 55% | 61.03°C |
| asrA_probe_25 | CCTTCACTTCGTCACTGGGAT | 21-mer | 52% | 62.29°C |
| asrA_probe_26 | TCCTTGAGGAAGAATTCGCG | 20-mer | 50% | 60.59°C |
| asrA_probe_27 | TCCTGCTGGATGATCTTCAAC | 21-mer | 47% | 60.29°C |
| asrA_probe_28 | ATCTCGCCCTTGAAACTACC | 20-mer | 50% | 59.94°C |
| asrA_probe_29 | CGACGCTGAAACGGTAGAAC | 20-mer | 55% | 61.81°C |
| asrA_probe_30 | ATCACCTCGACTTCCTTCAAC | 21-mer | 47% | 60.22°C |
| asrA_probe_31 | GTAGTGGTCGAGGAATTCGAC | 21-mer | 52% | 60.69°C |
| asrA_probe_32 | CCAGTTTTTCCTCGGAGATGTAG | 23-mer | 47% | 61.19°C |
| asrA_probe_33 | CGAGCTGTTTCTCCAACTGG | 20-mer | 55% | 61.43°C |
| asrA_probe_34 | CGATCTTCACCTTCGACTCG | 20-mer | 55% | 60.71°C |
| asrA_probe_35 | GATGTAGCTATAGGCGATCTCG | 22-mer | 50% | 60.20°C |
| asrA_probe_36 | GGTCGAAGAAGGTCGGGT | 18-mer | 61% | 61.46°C |
| asrA_probe_37 | GATCAGCTCGAAGATCTTCTGC | 22-mer | 50% | 61.36°C |
| asrA_probe_38 | GAGATAGTCGGGCAACTCTTC | 21-mer | 52% | 60.42°C |
| asrA_probe_39 | TAGTTCTTCAGTTCCTCGCTGTAC | 24-mer | 45% | 62.40°C |
| asrA_probe_40 | CGGCAAGACCTTCTCCAT | 18-mer | 55% | 59.46°C |
| asrA_probe_41 | GCGGACAGTTCCTTCTGCA | 19-mer | 57% | 63.01°C |
| asrA_probe_42 | ACTTCCACTTCCTTGCGCA | 19-mer | 52% | 62.60°C |
| asrA_probe_43 | GCTCTTGTCGTCCTTGGTGA | 20-mer | 55% | 62.59°C |
| asrA_probe_44 | AAATGCACGGTCAGCCCTT | 19-mer | 52% | 63.00°C |
| asrA_probe_45 | GCTTGTCCTTGCCGTACAC | 19-mer | 57% | 61.79°C |
| asrA_probe_46 | ATCACTTCCATCCGGTCGA | 19-mer | 52% | 60.86°C |
| asrA_probe_47 | GTCCTCGAGGTCCCTGG | 17-mer | 70% | 60.87°C |
| asrA_probe_48 | GCCTTTCTTCGGCACCTG | 18-mer | 61% | 61.51°C |
| pqsAB_probe_1 | GCGGAACAGAACCTCGGT | 18-mer | 61% | 62.46°C |
| pqsAB_probe_2 | CTCAGCGACGGTGCATC | 17-mer | 64% | 61.00°C |
| pqsAB_probe_3 | CAGCGAGGCATAGATGGC | 18-mer | 61% | 60.81°C |
| pqsAB_probe_4 | GCTGCTCAACAGCTCCCT | 18-mer | 61% | 62.50°C |
| pqsAB_probe_5 | CGGCCCAGAATTCGAACTC | 19-mer | 57% | 61.28°C |

| | | | | |
|---|---|---|---|---|
| pqsAB_probe_6 | CACCAACAGCACGCCTTG | 18-mer | 61% | 62.42°C |
| pqsAB_probe_7 | GCCAGTAACCCGGACTCAG | 19-mer | 63% | 62.49°C |
| pqsAB_probe_8 | GTACCAGCCACCTGCGAA | 18-mer | 61% | 62.48°C |
| pqsAB_probe_9 | CGGCAGATGACGGCAGAT | 18-mer | 61% | 62.69°C |
| pqsAB_probe_10 | GTTGGCCAATGTGGACATGA | 20-mer | 50% | 61.39°C |
| pqsAB_probe_11 | GCGGTATCGGGATCGAAATC | 20-mer | 55% | 61.21°C |
| pqsAB_probe_12 | GTCTGGCCCCGATAGTGATAA | 21-mer | 52% | 61.61°C |
| pqsAB_probe_13 | GAAGGCGAGTCGTTCAACG | 19-mer | 57% | 61.83°C |
| pqsAB_probe_14 | GGGACTTGGGATTGATCACG | 20-mer | 55% | 60.95°C |
| pqsAB_probe_15 | GGCAGTCGGCAGCGATAT | 18-mer | 61% | 62.76°C |
| pqsAB_probe_16 | GACCAGGTTCTCCAGAACCC | 20-mer | 60% | 62.34°C |
| pqsAB_probe_17 | CACTCATAGCCAGGCAACG | 19-mer | 57% | 61.28°C |
| pqsAB_probe_18 | CACAGTGACGGTAGGCACC | 19-mer | 63% | 63.06°C |
| pqsAB_probe_19 | CAGCACATGCAATTGGCTG | 19-mer | 52% | 60.91°C |
| pqsAB_probe_20 | TGGGAGAGAATGTAGGTCCG | 20-mer | 55% | 60.89°C |
| pqsAB_probe_21 | ATGGAAGGCACTCCAATCGA | 20-mer | 50% | 61.77°C |
| pqsAB_probe_22 | GAGGTGTATTGCAGGAAACAGG | 22-mer | 50% | 61.71°C |
| pqsAB_probe_23 | GCAGAAACCGAGCGTGTTG | 19-mer | 57% | 62.70°C |
| pqsAB_probe_24 | AGCAACTCCGTAGCGAACG | 19-mer | 57% | 63.09°C |
| pqsAB_probe_25 | AATCGAATACAGCCGGTCTC | 20-mer | 50% | 60.09°C |
| pqsAB_probe_26 | TGCCATAGCCGAAGAACATCT | 21-mer | 47% | 62.09°C |
| pqsAB_probe_27 | ACCAGGGAAAGAACAGGCTG | 20-mer | 55% | 62.57°C |
| pqsAB_probe_28 | TTCCTCGATAGTGTGTCCTTCG | 22-mer | 50% | 62.03°C |
| pqsAB_probe_29 | GGCCATTCACCTTGAACAGATC | 22-mer | 50% | 62.03°C |
| pqsAB_probe_30 | GGCAGGTAGGAACCAGAACC | 20-mer | 60% | 62.69°C |
| pqsAB_probe_31 | AACAGGGTCGGACGCAAG | 18-mer | 61% | 62.78°C |
| pqsAB_probe_32 | CATGTGCGAGGGAATCTGTT | 20-mer | 50% | 60.88°C |
| pqsAB_probe_33 | AAGGTTGTCGTGATAAAGGGTG | 22-mer | 45% | 61.06°C |
| pqsAB_probe_34 | GAAGACATGCGCGCGATTC | 19-mer | 57% | 62.95°C |
| pqsAB_probe_35 | CAGGCTCGAGTCGGTGAG | 18-mer | 66% | 62.27°C |
| pqsAB_probe_36 | GACCAGGACGTTGCGATAG | 19-mer | 57% | 60.69°C |
| pqsAB_probe_37 | TTGAGGTGTCCCTTGACGT | 19-mer | 52% | 61.25°C |
| pqsAB_probe_38 | CTTCGGGCAATCCAGGTTG | 19-mer | 57% | 61.52°C |
| pqsAB_probe_39 | CCAGACGGGTGAATCCGG | 18-mer | 66% | 62.62°C |
| pqsAB_probe_40 | CTCCAGACACACATAGGATGG | 21-mer | 52% | 60.08°C |
| pqsAB_probe_41 | ATCAGATGGTCGGGAGACAG | 20-mer | 55% | 61.25°C |
| pqsAB_probe_42 | AGGGTATCGACCACGTAGAG | 20-mer | 55% | 60.66°C |
| pqsAB_probe_43 | GCGAATATCGGTATTCAGCG | 20-mer | 50% | 59.47°C |
| pqsAB_probe_44 | CAGTTCAGTTGCGGGAAGTC | 20-mer | 55% | 61.71°C |
| pqsAB_probe_45 | GCGAGAAATCGTCGAGCAAAG | 21-mer | 52% | 62.70°C |
| pqsAB_probe_46 | AGCAGTTCATCCAGACGGT | 19-mer | 52% | 61.40°C |
| pqsAB_probe_47 | AGGATCTGGTTGTCGTCCA | 19-mer | 52% | 60.71°C |
| pqsAB_probe_48 | GAATCAACATGCCCGTTCCT | 20-mer | 50% | 61.17°C |

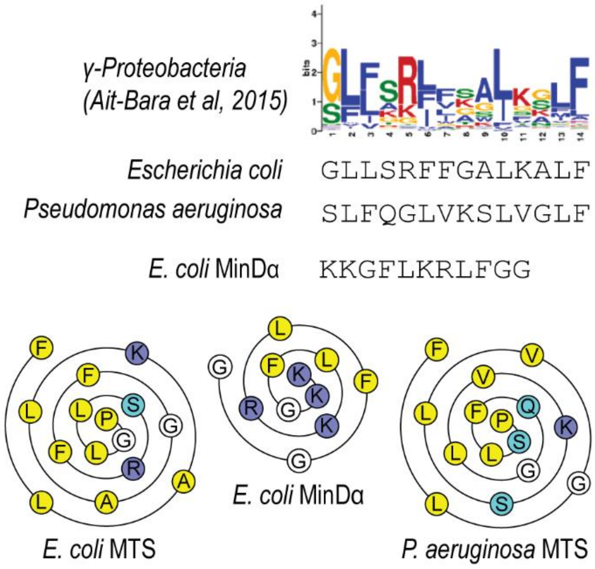

**Appendix Figure S1:** Amino acid sequence and helical wheel projection of the RNase E amphipathic helix (MTS) from *E. coli* or *P. aeruginosa* compared to the unrelated amphipathic helix from the well-characterised *E. coli* MinD amphipathic helix (MinDα) (Szeto *et al*, 2003; Zhou & Lutkenhaus, 2003). The RNase E MTS conservation logo plot from (Aït-Bara *et al*, 2015) is included for reference.

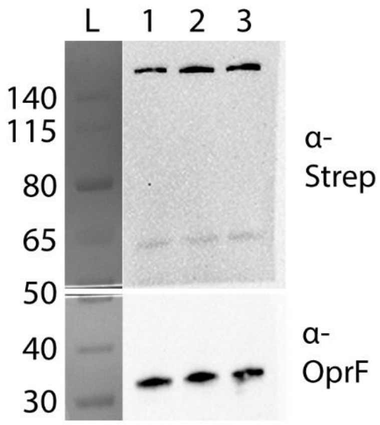

**Appendix Figure S2:** Western Blot of crude lysates of strains expressing RNase E-Strep, RNase $E^{\Delta MTS}$-Strep, or RNase $E^{MinD\alpha}$-Strep (chromosomal expression from the native locus). Protein levels are unaffected by the deletion of the MTS or the replacement of the MTS by the MinD amphipathic helix. 1: RNase E-Strep; 2: RNase $E^{\Delta MTS}$-Strep; 3: RNase $E^{MinD\alpha}$-Strep. Western Blot against the constitutive OprF protein was performed as a loading control.

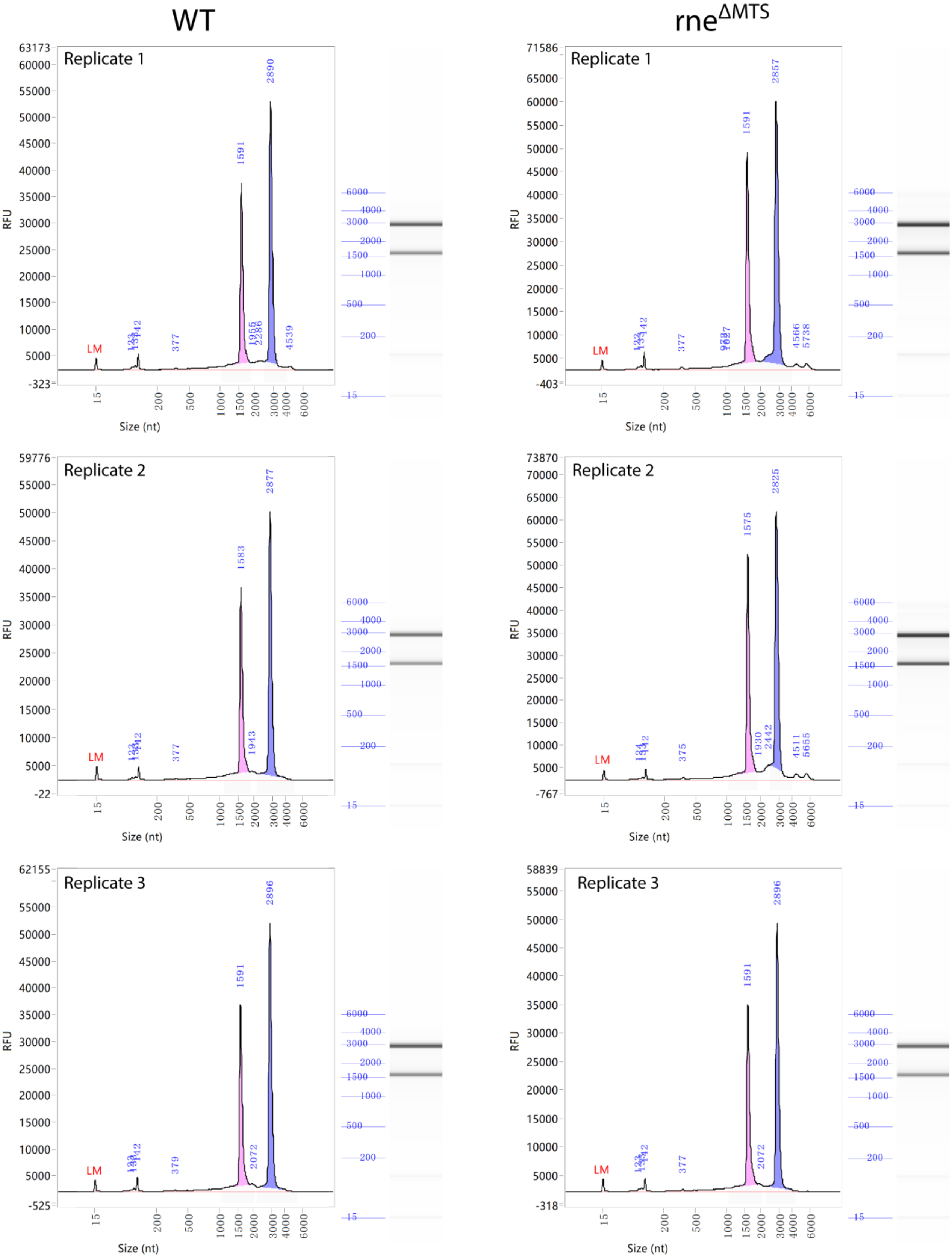

**Appendix Figure S3**: Agilent 5400 peak profiles and simulated electrophoresis of total RNA samples of the WT or *rne*ΔMTS strains (biological triplicates). The profile of each replicate is shown for each strain. Analysis was performed by Novogene prior to RNA sequencing.

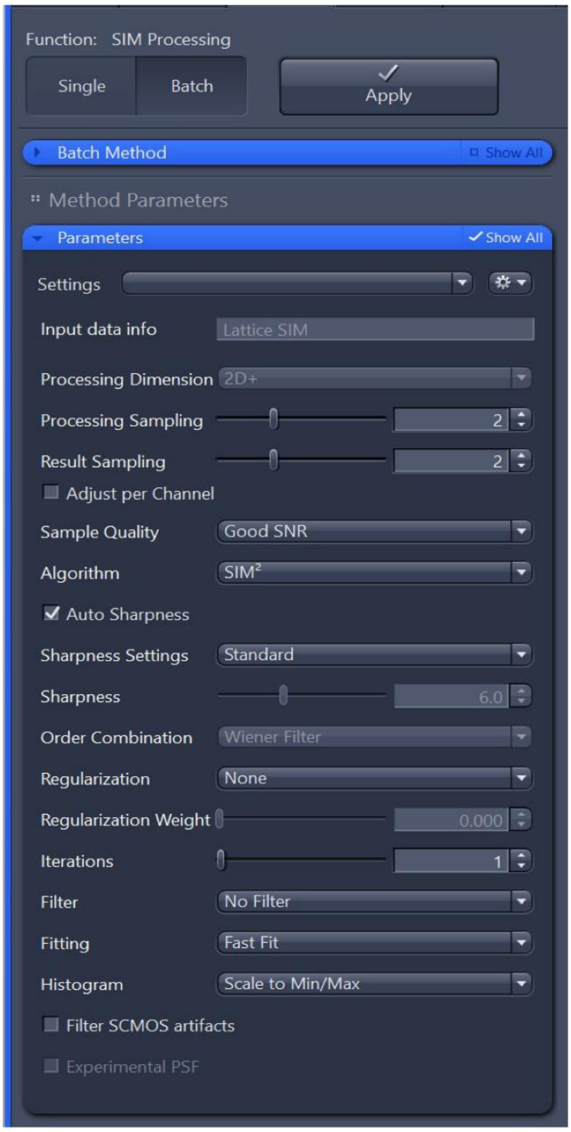

**Appendix Figure S4:** Reconstruction parameters used for SIM images in this study. The Zen Blue software was used for image acquisition and processing.

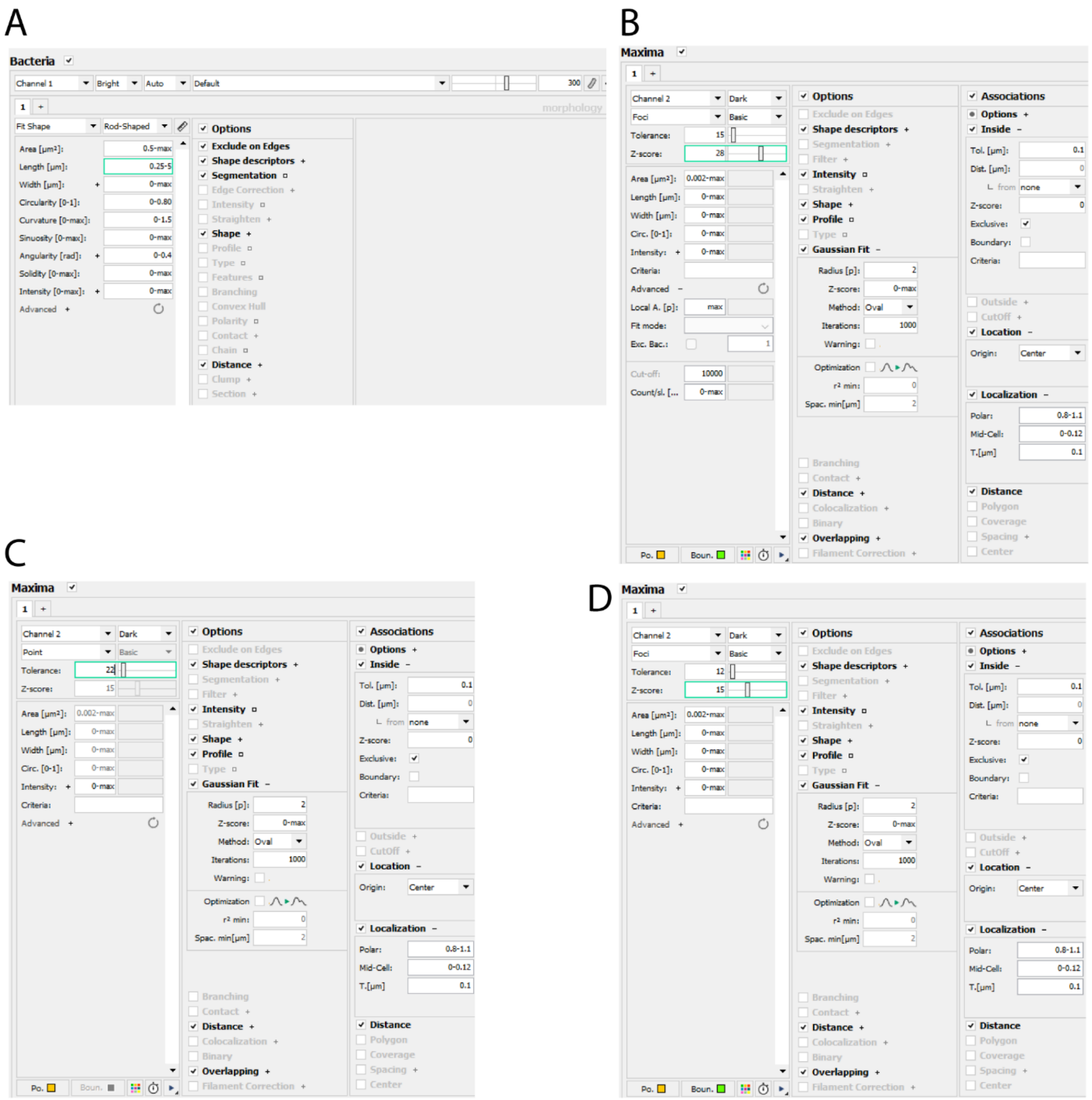

**Appendix Figure S5**: MicrobeJ detection parameters used in this study. Parameters that were manually adjusted between different images are highlighted by a green box. **(A)** Settings used for bacteria detection in all analysis done. The parameter of cell length (green box) was sometimes decreased to a max value of 4.5 or increased to a max value of 5.5 to ensure good cell detection in each image. When needed, additional manual segmentation was done as described in Materials and Methods. **(B)** Settings used for maxima detection when analysing msfGFP-tagged RNase E variants. A foci detection mode was chosen (using Z-score) to allow characterisation of foci morphology and subcellular localisation. The tolerance value was set to 15 for all image analysis but the Z-score was adjusted for each image to ensure optimal detection of foci outlines based on visual inspection. Z-score values set ranged from 26 to 58. **(C)** Settings used for maxima detection to obtain foci counts and intensities of mRNAs visualised by FISH. A point detection mode was chosen (no Z-score) to simplify the analysis and avoid manual input. The tolerance value was set identically for all images analysed for a given mRNA. For putP, asrA, and pqsAB, the tolerance values were 20, 22, and 28, respectively. **(D)** Settings used for maxima detection to plot subcellular localisation of mRNAs visualised by FISH. A foci detection mode was chosen (using Z-score) to ensure very low rate of false positives (at the cost of more false negatives) and very accurate detection of foci centre positioning thanks to detected foci outlines. The tolerance value was set to 12 for all image analysis but the Z-score was adjusted for each image to ensure optimal detection of foci outlines based on visual inspection. Z-score values set ranged from 7 to 15.

#### Appendix Supplementary Methods

##### Twitching and swimming assays

The experimental conditions and procedures were essentially as described in (Valentini *et al*, 2016), with minor changes: 1% LB Lennox was used for the twitching assays, and both swimming and twitching plates were incubated at 37°C.

Aït-Bara S, Carpousis AJ, Quentin Y (2015) RNase E in the γ-Proteobacteria: conservation of intrinsically disordered noncatalytic region and molecular evolution of microdomains. *Mol Genet Genomics* 290: 847-862

Marshall OJ (2004) PerlPrimer: cross-platform, graphical primer design for standard, bisulphite and real-time PCR. *Bioinformatics* 20: 2471-2472

Szeto TH, Rowland SL, Habrukowich CL, King GF (2003) The MinD Membrane Targeting Sequence Is a Transplantable Lipid-binding Helix*. *Journal of Biological Chemistry* 278: 40050-40056

Valentini M, Laventie B-J, Moscoso J, Jenal U, Filloux A (2016) The Diguanylate Cyclase HsbD Intersects with the HptB Regulatory Cascade to Control Pseudomonas aeruginosa Biofilm and Motility. *PLOS Genetics* 12: e1006354

Zhou H, Lutkenhaus J (2003) Membrane Binding by MinD Involves Insertion of Hydrophobic Residues within the C-Terminal Amphipathic Helix into the Bilayer. *Journal of Bacteriology* 185: 4326-4335
